# Emergence of growth and dormancy from a kinetic model of the *Escherichia coli* central carbon metabolism

**DOI:** 10.1101/2021.07.21.453212

**Authors:** Yusuke Himeoka, Namiko Mitarai

## Abstract

Physiological states of bacterial cells exhibit a wide spectrum of timescale. Under nutrient-rich conditions, most of the cells in an isogenic bacterial population grow at certain rates, while a small subpopulation sometimes falls into a dormant state where the growth rates slow down by orders of magnitude. The dormant cells have unique characteristics: The metabolic activity is quite slow, and the dormant cells typically exhibit a high tolerance for a range of stresses, such as antibiotics applications. To reveal the origins of such heterogeneity of timescales, we constructed a kinetic model of *Escherichia coli* central carbon metabolism, including the dynamics of the energy currency molecules, and asked if perturbations of the metabolites’ concentrations lead to the distinct metabolic states. By numerically studying the relaxation dynamics, we found that the model robustly exhibits two qualitatively distinct relaxation dynamics depending on the initial conditions generated by the perturbations. In the first type, the concentrations of metabolites reach the steady-state quickly, resembling the growing dynamics. On the other hand, the other type of dynamics takes a much longer time to reach the steady-state, and during the relaxation, cell growth almost halts, reminding us of the dormant cells. In order to unveil the mechanism of distinct behaviors, we reduced the metabolic network model into a minimal model without losing the emergence of distinct dynamics. Analytical and numerical studies of the 2-variable minimal model revealed the necessary conditions for the distinct behavior, namely, the depletion of energy due to the futile cycle and its non-uniform impact on the kinetics because of the coexistence of the energy currency-coupled and uncoupled reactions as well as branching of the network. The result is consistent with the experimental reports that the dormant cells commonly exhibit low ATP levels and provides a possible explanation for the appearance of dormant cells that causes antibiotic persistence.

## Introduction

Bacterial growth rates span a wide range of timescales: *Escherichia coli* cells typically double every 20 minutes under nutrient-rich conditions, while cells can also exhibit dormancy where the growth of cells almost halts and yet the death is strongly suppressed [1–3]. The transition to the dormant states can either be a stochastic event or a response to hostile environments such as starvation and exposure to antibiotics. This dormancy is a beneficial strategy for surviving nutrient-poor conditions as it can lower the cell’s nutrient requirements [4]. Also, dormancy is known as the main cause of bacterial persistence that has a high tolerance to antibiotics, and thus, has been gathering attention from a wide range of fields from microbiology to therapeutic studies [2,3,5,6].

Notable changes in the timescale of cellular physiology are happening in the dormant cells. It has been implied that the dormant cells have a sort of memory capacities: The lag time was shown to depend on the length of time that the cells are starved [7–9] and the death rates of the starved cells differ depending on the previous culture conditions even though the starvation condition is identical [10]. Given that slow dynamics are vital for storing memories, a drastic change in the timescale of cellular physiology is necessary. Indeed, it was reported that the proteome kept changing at least for 8 hours in the starved *E. coli* cells [11].

Experimental studies have revealed the links between dormancy and several molecules, such as growth-inhibiting genes and metabolic enzymes (cf. reviews of Refs. [3,12,13]). Based on the experimental findings, models that exhibit the transition to dormancy have been proposed; The model developed by Klumpp *et al.* [14] shows the bistability of growing- and dormant state led by the toxin-based feedback mechanism of gene expression. The transition mechanism suggested by Rocco *et al.* is based on the bursting activity of gene expression [15,16]. According to the model by Radzikowski *et al.* [11], the collapse of the metabolic homeostasis by perturbation is the key to the transition: *A* strong perturbation is applied to the metabolic state, and the resulting low metabolic flux cannot support the synthesis of the metabolic enzymes to restore metabolic homeostasis. This failure of metabolic re-adjustment further lowers metabolic activity.

All the models mentioned above for exhibiting the dormancy transition include the gene expression dynamics. Here in the present manuscript, we explored another possibility: the dormancy transition is triggered by the metabolic dynamics itself, without regulatory changes. Metabolic reaction networks are highly interconnected via cofactors such as ATPs; thus, the kinetic models of metabolic networks should have high non-linearities. The emergence of different timescales is one of the hallmarks of nonlinear dynamical systems. Indeed, the studies of a simple catalytic reaction network showed that the relaxations to the steady-state are much slower than that inferred from the rate constants of the reactions and exhibit multiple plateaux [17–19].

In the present manuscript, we study the kinetic model of *E. coli* central carbon metabolism with cofactors, such as ATP, as variables. To the best of our knowledge, this is the first kinetic model of *E. coli* metabolism where the dynamics of cofactors are dealt as variables: There are a number of studies of kinetic model of *E. coli* central carbon metabolism [20–35]. However, as far as we know, the dynamics of the cofactors are neglected [20–30], or if included, the relaxation dynamics of the models are not actually computed [31–35]. As the experimental studies suggested [36,37], ATPs may play a central role in the transition to dormancy. Thus, the cofactors can be vital model components for studying the growth-dormancy transition.

In the following sections, we present that the kinetic model of *E. coli* central carbon metabolism with cofactors robustly exhibits two distinct dynamics: One is reminiscent of the normal growth behavior, and the other is analogous to the dormant dynamics. Then we derive the minimal network showing qualitatively the same dynamics. The minimal model analysis reveals two necessary conditions for the emergence of both growth and dormant dynamics: the depletion of energy due to the futile cycle and its non-uniform impact on the kinetics because of the coexistence of the energy currency-coupled and uncoupled reactions as well as branching of the network.

The obtained result implies that the depletion of ATP and ADP itself leads to the slow dynamics of the metabolites’ concentrations. This conclusion is consistent with the “low-energy” view of the bacterial dormancy presented in [36,38], and highlights the notable impact of introducing cofactors into models. We also discuss the possible applications of our analysis for the studies of dormancy in other species based on the minimal network motifs.

## Results

### Model

In the present manuscript, we study the *E. coli* core network [39] as one of the simplest models of the real metabolic reaction networks. The *E. coli* core model was obtained from BiGG database [40]. The model contains stoichiometry and the reversibility of the reactions. The *E. coli* core model has 52 and 75 intracellular metabolites and reactions, respectively. After an appropriate data curation as described later, we implemented the model by using the ordinary differential equation (ODE) that describes the dynamics of concentrations of metabolites.

We applied several modifications to the model to make it suitable for ODE implementation. First, small molecules such as O_2_, H_2_O, and NH_4_, were not considered as variables but treated as constants under the assumptions that the external concentration of these chemicals are kept constant, and uptakes/secretions of them take place quickly. The uptake and secretion pathways of all carbon sources except glucose are removed.

Under anaerobic conditions, cells transfer the free energy to ATP directly, while under aerobic conditions, most of the energy transfer takes an indirect form: the energy is first transferred to other chemicals such as NADH and NADPH, and then, the stored energy in NADH, NADPH, and others are used for converting ADP to ATP. The conversion yield of ATP per NADH in the *E. coli* core model is 1.25 (via NADH16, CYTB, and ATPS4r), and NADH/NADPH yield is roughly unity. For introducing the cofactors to the model in a simple manner, we assume that the energy transfers via NADH and NADPH are sufficiently fast and ATP/NADH(NAPDH) yield as unity. According to these assumptions, we replace NAD(NADP) and NADH(NADPH) with ADP and ATP, respectively (we discussed the validity of this assumption in the Discussion section and *SI text* Section.6). Full lists of the chemical components and the reactions are provided in *SI Data.* 1.

Also, the stoichiometry of the growth reaction was modified. The original *E. coli* core model has the biomass production reaction leading to the cell growth consisting of 16 substrates and 7 products with non-integer stoichiometry constants. For kinetic modeling, such reactions having too many substrates and products lead to numerical instability, and non-integer stoichiometry is unreasonable. Thus, we replaced the biomass production reaction with a following reaction: (Erythrose 4-phosphate) + (L-Glutamine) + (ATP) → (ADP). This reaction is much simpler than the original one. Still, it requires the model to run all the modules of the metabolic reactions, namely the pentose phosphate pathway for Erythrose 4-phosphate (e4p), TCA cycle for L-Glutamine (gln), and energy generation for ATP. Hereafter, we call this simplified biomass production reaction as the growth reaction.

The resulting model consists of 32 variables and 40 reactions. The final metabolic reaction network is drawn in Fig. 1. Our model cell takes up the nutrient from the node labeled as “glc” which has a constant concentration, performs successive conversion of the chemicals generating energy, and proceeds with the growth reaction.

**Figure 1:**
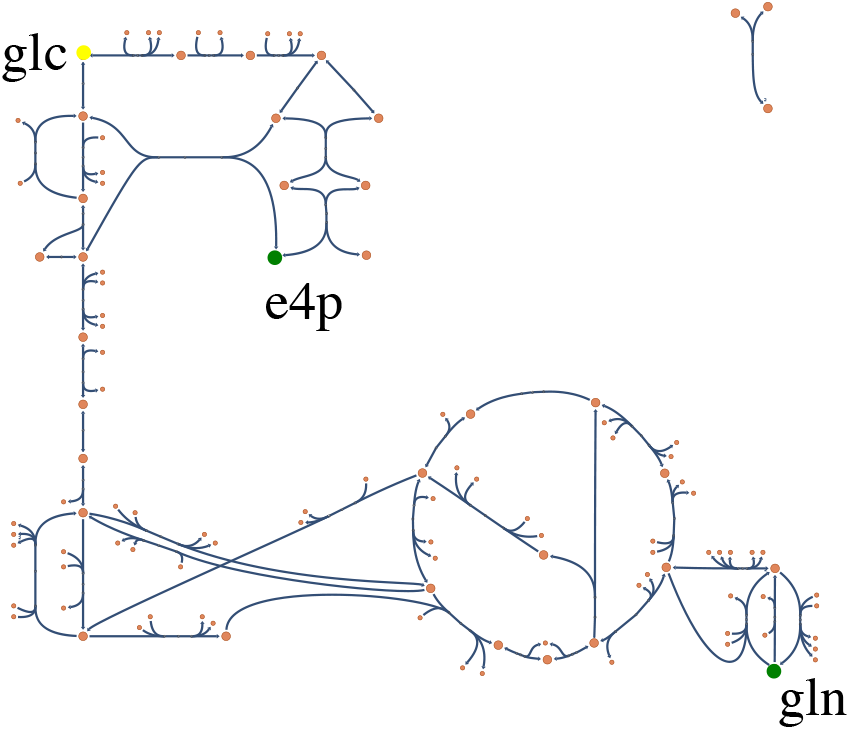
The metabolic network of the *E. coli* core model generated by Escher [40]. The sole carbon source (glucose-6-phosphate) is placed at the left top (abbreviated as glc). We highlighted the substrates of the growth reaction other than ATP, namely, e4p and gln. The growth reaction is not drawn.

First, we simulated the model with realistic setups. The kinetic parameters of *E. coli* core model have been estimated using the metabolic ensemble modeling (MEM) by Khodayari and colleagues [33]. We derived the Michaelis-Menten type rate equation for each reaction according to the enzyme kinetics used in [33] with the presented kinetic parameters. Then we assumed that each chemical species is consumed/synthesized by associated reactions, diluted as the cell grows, and spontaneously degraded at a slow rate. Thus, the temporal change of the concentration of the ith chemical species *X_i_* is ruled by

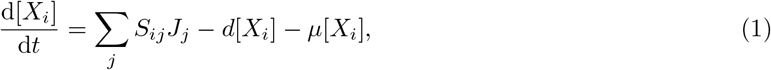

where *S* is the stoichiometric matrix, and *J_i_*s are the fluxes due to chemical reactions. *d* and *μ* are the spontaneous degradation rate and the growth rate, respectively. Note that the concentrations of enzymes are supposed to be constant and lumped in the kinetic parameters. We assumed that spontaneous degradation is a very slow process represented by a single parameter. The dilution and degradation terms are omitted in the AMP, ADP, and ATP equations because the de-novo synthesis of the adenine nucleotide carriers is not modeled in the *E. coli* core model. This assumption is equivalent to the homeostasis of the total adenine nucleotide carriers. (We check that the assumption can be relaxed by introducing a phenomenological reaction for the de-novo synthesis of AMP, see *SI Text* Section.5). According to the growth reaction which we have introduced above, our model cell grows as the reaction (Erythrose 4-phosphate) + (L-Glutamine) + (ATP) → (ADP) proceeds. We chose the simplest kinetics of the growth reaction given by *J_g_* = *v_g_* [e4p][gln][atp] and the growth rate as *μ* = *rJ_g_*. We fit the values *v_g_* and *r* so that the growth rate at the steady-state is in the range of the typical growth rate of *E. coli* in minimal glucose media ≈ 0.5 per hour. The spontaneous degradation rate *d* is set to be one-hundredth of the steady growth rate so that the effect of the spontaneous degradation is negligible at the attractor. The concentration of the nutrient ([glc]) and the total concentration of the adenine nucleotide carriers (*A_t_*) are set to 20mM and 1mM, respectively (see *SI Text* Section.1).

To see how many attractors the model has, we computed the model dynamics from multiple initial concentrations. As far as we have checked, the model has only a single steady-state attractor.

### Dormant trajectory

We applied random perturbations to the steady-state concentration to emulate the exposure to the sub-lethal stresses that disturb intracellular states. The perturbations are applied in a multiplicative manner. For the metabolite *i* with the steady-state concentration 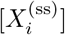, the initial concentration is given by 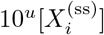, where *u* is a random number sampled from uniform distribution in [−2, 2]. The concentrations of ATP, ADP, and AMP are normalized so that the total concentration is *A_t_* because it is conserved in the model.

We found that the model exhibited two qualitatively distinct relaxation behaviors depending on the initial conditions. The typical time course of each type is plotted in Figs. 2 A in log scale to depict the wide range of the concentration and timescale.^1^

**Figure 2:**
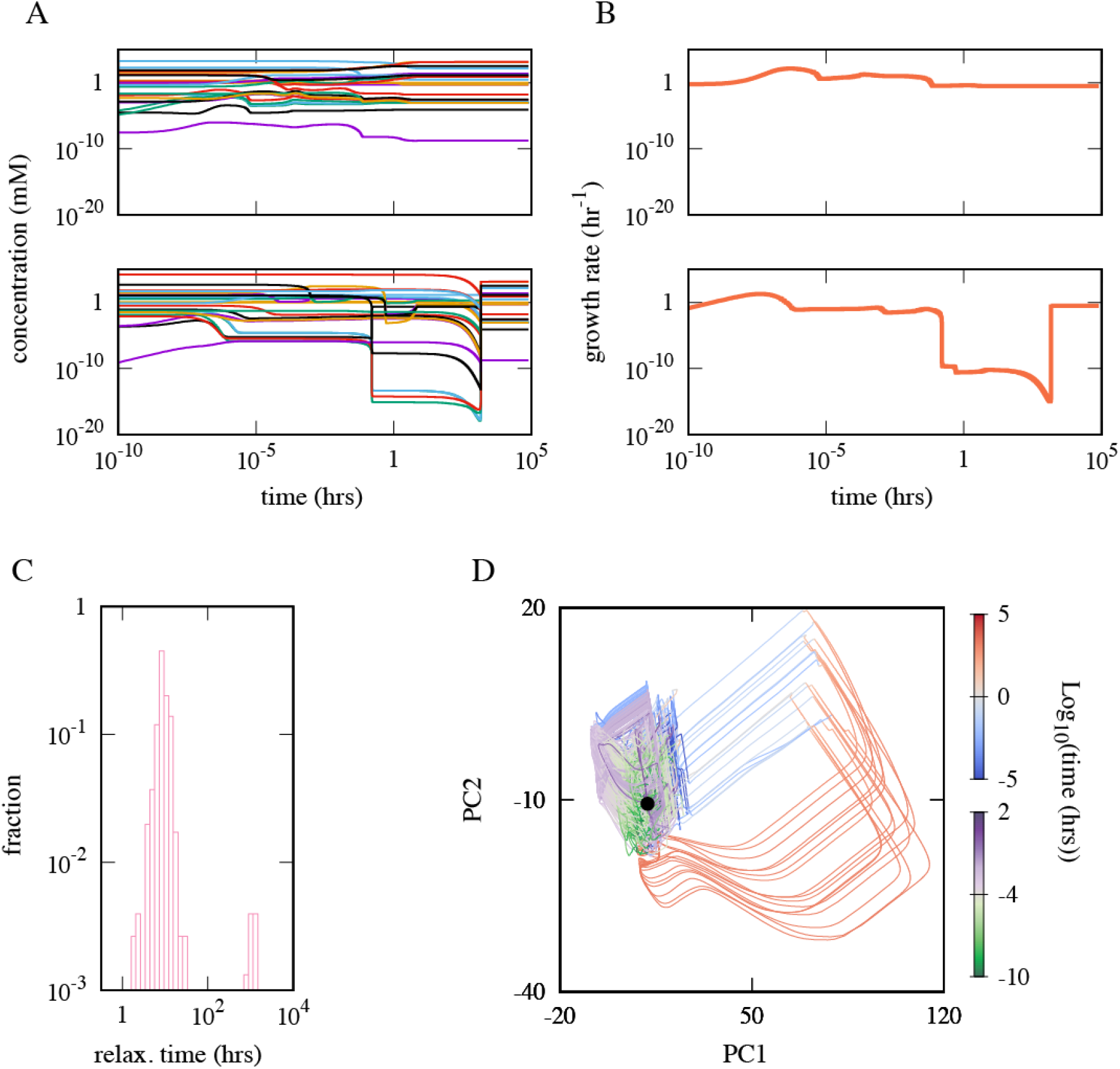
A. Two characteristic dynamics of *E. coli* core model starting from different initial points. While the growth rate of the cell is ≈ 0.5 per hour at the attractor, there are huge differences in the relaxation behaviors between the top and bottom panels. B. The temporal changes of the growth rates along the dynamics in the same row in panel A are plotted. C. The distribution of the relaxation time shows a clear bimodality. C. Trajectories are overlaid in 2-dimensional principal component space. The color indicates log_10_ of time. The trajectories having shorter relaxation time (several hours) are colored green-white-purple while the others are colored blue-white-red. The black point corresponds to the steady-state attractor. Initial concentration of each metabolites is 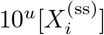 (mM) with 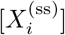 as the steady-state concentration of the ith metabolite, and *u* as a random number uniformly distributed in [−2, 2] while the total concentrations of adenine nucleotide carriers are normalized. Parameters other than ones obtained from [33] are [glc] = 20(mM), *A_t_* = 1(mM), *d* =5 × 10^−3^(hour^−1^), *v_g_* = 3.6 × 10^4^(mM^−2^· sec^−1^) and *r* = 5.0(mM^−1^).

Even though the two trajectories eventually relax to the same steady state, the relaxation behaviors are evidently distinct. First, the concentrations after minutes (~ 10^−1^ hours) are different between the top and bottom panels by many orders of magnitude. The concentrations of several chemicals are smaller than one molecule per cell, especially in the bottom panel. We revisit this point in the discussion section. Also, the characteristic timescale between them is clearly different. The concentrations of the chemicals reach close to the steady values in minutes in the top panel.

In contrast, the concentrations keep changing for a much longer time, *t* ≈ 10^3^ hours in the bottom panel, which is experimentally indistinguishable from the situation where cells stop growing. When sampled over various initial conditions, the relaxation time distribution has a clear bimodality as shown in Fig. 2C. Here, the relaxation time is defined as when the distance between the steady-state attractor and the state in the logarithm-converted phase space first becomes less than 0.05.

For visualizing the differences among the trajectories, we analyzed all the trajectories in the phase space by the principal component analysis (PCA, see Materials and Methods), where all the trajectories are converted to the logarithmic scale. We plotted all trajectories projected onto the 2-dimensional principal component space (PCS) in Fig. 2D. The trajectories were classified into two groups by the relaxation time and differently colored. The first group is quickly-relaxing trajectories that the trajectory in the top panel of Fig. 2A belongs to (colored in green-white-purple). The trajectory in the bottom panel of Fig. 2A is grouped into the other group, colored blue-white-red, which takes much longer to relax to the steady-state attractor.

The remarkable gaps between the timescale of chemical reactions and, accordingly, the growth rate during their relaxations highlight the difference between the two time courses. The specific growth rate *μ* at the steady-state is ≈ 0.5 hour^−1^, and the model cell achieves this growth rate in a few seconds in the top panel of Figs. 2A, while less than 10^−10^ hour^−1^ in the bottom panel at *t* = 10^2^ hours (at plateau). Thus, in the following sections, we call the trajectories of the second group “dormant trajectories” because of their much slower growth rate than the other group. Accordingly, the trajectories of the first group are termed “growth trajectories”. The following sections are devoted to unveiling the mechanism leading to the differentiation of the growth and dormant trajectories.

### Systematic Model Reduction

In the previous section, we saw that distinct relaxation dynamics emerge depending on the initial concentrations of the metabolites. Interestingly, we found that the emergence of distinct dynamics is a robust feature of the *E. coli* core model. The distinct dynamics emerge even if we use the mass-action kinetics instead of the Michaelis-Menten function for the reaction-rate function. Also, it is less sensitive to the specific choice of the parameter values (see *SI Text* Section.2 and 8). This robustness implies that the distinct dynamics emerge from the structure of the metabolic reaction network of the *E. coli* core model rather than choices of specific parameter values.

Thus, it is worth asking if there are understandably simple, minimal network architecture(s) in the *E. coli* core network which lead to the distinct trajectories. In the present section, we reduce the *E. coli* core network to obtain a minimal network exhibiting distinct relaxation dynamics. As far as we know, there is no method to reduce the reaction network without losing the characteristic nature of the relaxation dynamics. Once the concentrations of the cofactors are dealt with as variables, metabolic reactions in the model get highly interconnected, and the well-known reduction method works poorly. For instance, adiabatic elimination may eliminate merely one or few reactions, and obtaining an understandably simple model is hopeless. Thus, here we focus only on the emergence of distinct trajectories. As will be seen, this allows us to derive a much simpler model than the original model.

In the following, we remove one or a few reactions from the network step by step and check if the reduced model still exhibits distinct trajectories (a solid criterion is introduced later). As illustrated in Fig. 3A, we consider two types of reaction removal, namely, simple removal and contraction. First, we describe the simple removal. Suppose that there are reactions *A ⇌ B, B ⇌ C* and *C ⇌ A*, and also A and *B* are connected to the rest part of the network by the other reactions (Fig. 3A-(i)). The simple removal removes the reaction *B ⇌ C* and *C ⇌ A*, and, accordingly, eliminates the chemical C because it is a disconnected component in the network.

**Figure 3:**
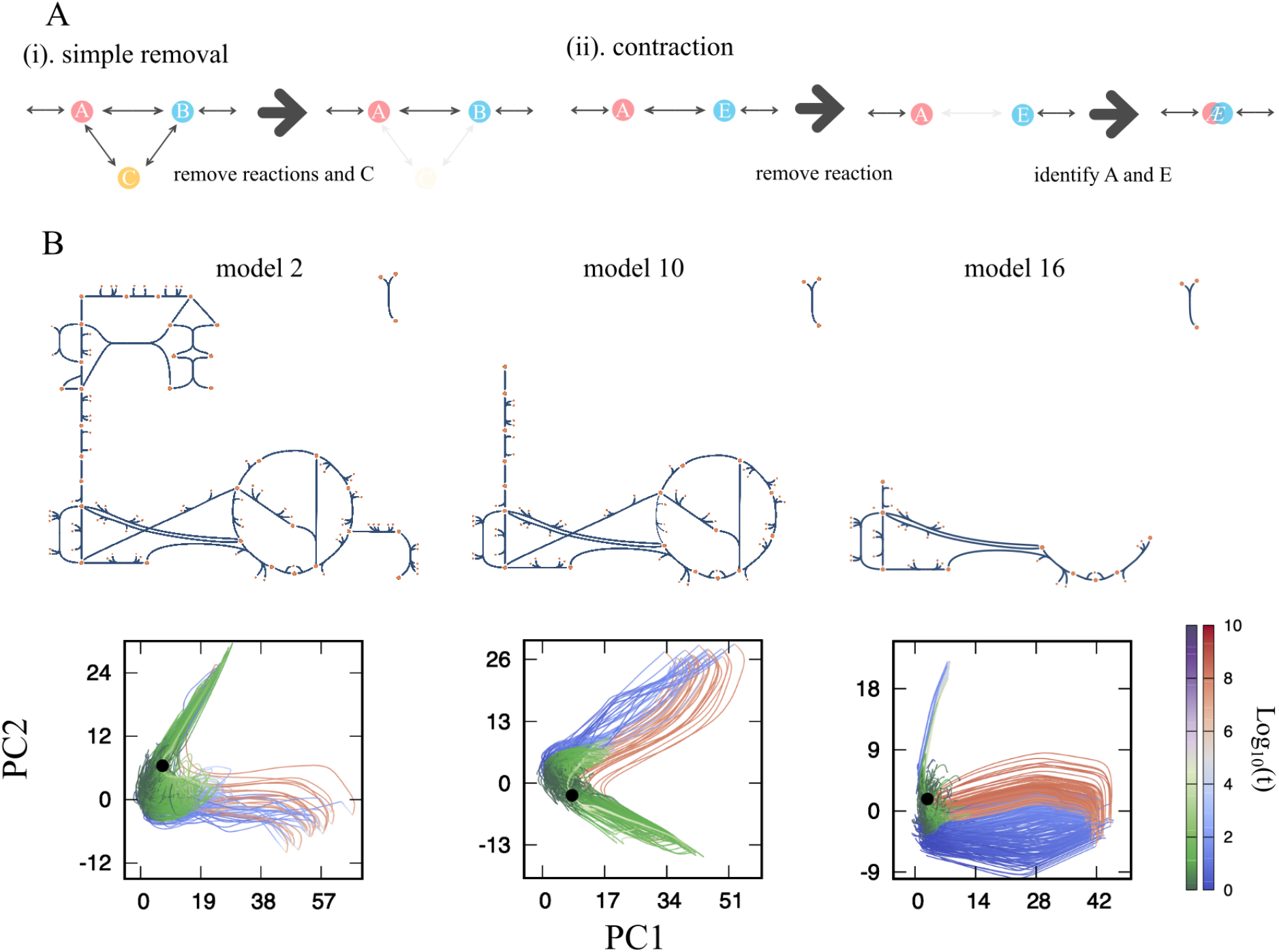
A. Two types of reaction removal. The simple removal removes one or a few reactions from the network. The number of reactions to be removed is determined so that the removal does not make the dead-end chemicals. The contraction removes a single reaction first, and then, the substrate and product of the removed reaction are identified and regarded as a new chemical species. B. The reduced networks of the intermediate models (models 2, 10, and 16) are drawn. The only nutrient (glucose) is at the top left corner of the network. The trajectories projected onto the PCS of each model are also shown. For the coloring protocol of the trajectories, see the main text.

In contrast, chemical species are merged by the contraction (Fig. 3A-(ii)). It removes a reaction *A ⇌ E*, and then, the chemical *A* and *E* are identified, forming a new chemical Æ. Here, we avoided the appearance of the dead-end chemical, which has only one reaction because networks with dead-end chemicals can cause a heavy accumulation of the chemicals, potentially leading to an artifactual anomalous relaxation behavior.

At each reduction step, we checked if the reduced model exhibits the two distinct classes of trajectories by computing its dynamics (For the details and the criterion for the distinct dynamics, see *SI Text* Section.3). In the following we use the model with the mass-action kinetics with most parameters to be unity because we have confirmed the distinct dynamics also emerge with this setup (for the detailed setups, see *SI Text* Section.2).

We have reduced the *E. coli* core model step-by-step according to the model reduction method described above (The pseudo-code is presented in Algorithm.S1 in the SI Text.). For accomplishing the network reduction, we manually determined the order of the reaction removal so that subsystems of the network are removed or contracted in consecutive reduction steps. We completed the model reduction by removing and contracting the L-glutamine synthesis pathway (4 steps), pentose-phosphate pathway (4 steps), glycolytic pathway (3 steps), and TCA cycle (7 steps) with the indicated number of steps in the parenthesis. The full list of the removed reactions is provided in *SI Data.1.* Note that we also tried the model reduction in random orders of the reaction removal (see *SI Text* Section.7). The minimal networks led by the reduction surely depend on the order of the reaction removal. However, all the minimal networks commonly satisfied the two conditions for the emergence of the distinct trajectories discussed later. We revisit the case of random-order reduction in the Discussion section.

The reaction network, and the trajectories projected onto the PCS of selected models are shown in Figs. 3C. We colored the trajectories based on the relaxation time of each. The figure shows that dormant trajectories (blue-white-red trajectories) commonly take detours to reach the attractor in the PCS. We confirmed that the dormant trajectory also takes detours in the original high-dimensional phase space (*SI Text* Section.3.5).

### A Minimal Model

After the 18 steps of reductions, we reached the stage where no more reduction is possible without losing the distinct dynamics. The reaction network and names that remained in this minimal network (model 18) are depicted together with the original *E. coli* core network in Fig. 4A. The network consists of glucose (glc), phosphoenolpyruvate (pep), pyruvate (pyr), oxaloacetate (oaa), ATP, ADP, and AMP. As highlighted in the original network, the reaction from glc to pep is the contraction of the glycolytic pathway, and oaa is representative of the chemicals in the TCA cycle. It is worth noting that the network’s local structure among pep, pyr, and oaa is unchanged (cyan boxes). In other words, the minimal network is obtained by removing the pentose phosphate pathway and contracting the glycolytic pathway and the TCA cycle. Also, the reaction ADK1 converting two ADPs to ATP and AMP is conserved. As shown in Fig. 4B and C, the model still exhibits distinct trajectories. In the following, we use one-letter variables instead of the abbreviations of the metabolites. *x,y, z* and *g* denote [pep], [pyr], [oaa], and [glc], respectively. *a,b* and *c* are assigned for [atp], [adp] and [amp], respectively. We use the upper-case characters for referring to the name of the metabolites (i.e., *X* indicates phosphoenolpyruvate, and *x* means its concentration [pep].)

**Figure 4:**
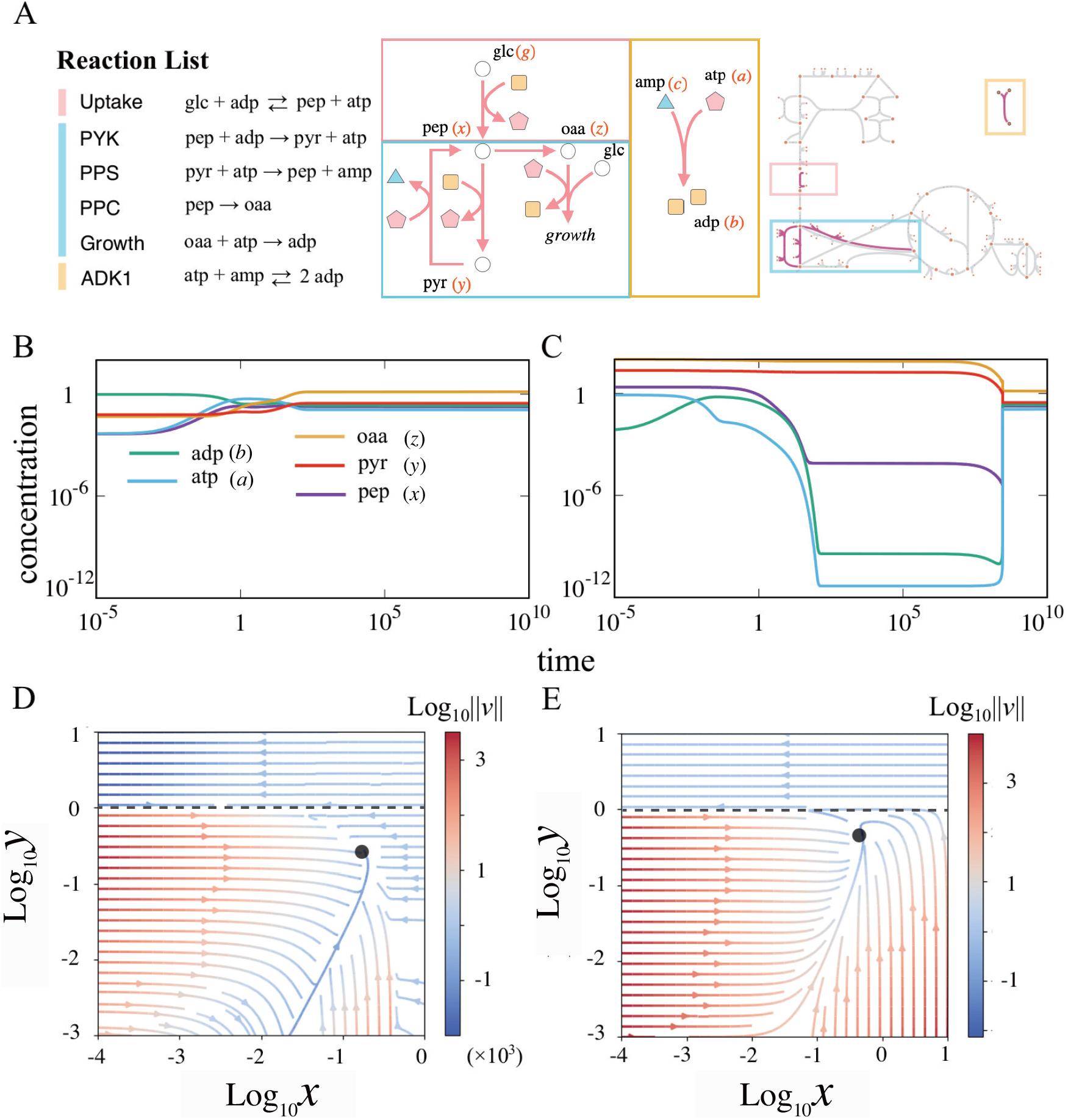
A. The list of the reactions in the minimal network (left). The structure of the minimal network (middle). The original network with the reactions in the minimal model is highlighted (right). The minimal model consists of three parts, namely, the glycolytic pathway (pink bar and boxes), the joint part between the glycolytic pathway and the TCA cycle (cyan bar and boxes), and the adenosine kinase reaction (yellow bar and boxes). B and C. Example time courses of the growth trajectory (B) and the dormant trajectory (C) of the minimal model. D. The streamline representation of the vector field of the 2-variable minimal model where the steady values of [atp], [adp] and [oaa] under given [pep] and [pyr] are numerically solved. E. The streamline representation of the simplified minimal model. The color indicates the norm of the vector *v* = (d*x*/d*t*, d[pyr]/d*t*) at each point, and the black dots indicate the attractor of each in D and E. *ϕ*_0_ = 10^−8^ in E. The dashed lines in D and E represent the boundary where the direction of the vector field changes dramatically (*y* = 1). The lines are drawn according to the Eq. (4) for D and the definition of *ϕ* for E.

The model consists of five variables (recall that *g* and *a* + *b* + *c* are constant). We wish to simplify the model further and draw the 2-dimensional vector field. Thus, we tried the adiabatic elimination of the concentrations of three chemicals (because now we have the five variables). By checking all the possible combinations of three chemicals, we found that the adiabatic elimination of *a, b*, and *z* does not lose the distinct dynamics. The set of ODEs of the minimal model is then given by

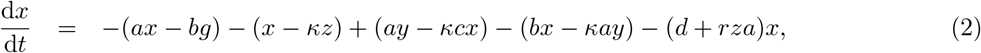

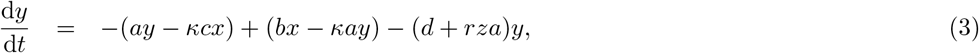

where *a, b* (accordingly *c* = *A_t_* – *a* – *b* with *A_t_* as the total concentration of adenosine energy carriers), and *z* are adiabatically solved, and thus, are the functions of *x* and *y*.

The two-variable system allows us to visualize the vector field. As shown in Fig. 4D, interestingly, there is a boundary below and above which the streamlines change the direction dramatically (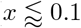 and *y* ≈ 1). Below the boundary, the state relaxes to the attractor rather straightforwardly, corresponding to the dynamics shown in Fig. 4B (growth trajectory). On the other hand, trajectories starting from the upper region first travel to the left side of the phase space (small *x* region) and return to the attractor, corresponding to Fig. 4C (dormant trajectory). We attribute the emergence of the distinct trajectories to this dramatic change in the direction of the vector field across the boundary. Hereafter, we call the region above- and below the boundary the dormant and the growth region, respectively.

### Conditions for the emergence of distinct trajectories

What determines the boundary between the growth- and the dormant region, and why is the vector field of the 2-variable model (Fig. 4D) almost parallel to the horizontal axis in the dormant region? For the first point, we found a large gap between the two regions in *a* and *b*. In the dormant region, their concentrations are low (~ 10^−8^), while in the growth region, they are in the order of 0.1. This gap gives an insight into the second point. We found that the drastic change in the direction of the vector field occurs because four of the five reactions of the model coupled with the adenine nucleotide carriers. Thus, these reactions halt almost entirely in the dormant region.

Intuitively, the low values of *a* and *b* in the dormant region can be understood from the reactions in Fig. 4A as follows. First, let us consider the situation where *x* and *y* are in the dormant region. If in addition, *x* is low, the uptake reaction proceeds in the direction *G* + *B* → *X* + *A*. When this reaction accumulates some ATP, PPS proceeds in the direction *Y* + *A* → *X* + *C*, because x is low. In total, the uptake reaction and PPS form a futile cycle that converts *A* and *B* into *C*. Note that the PPS does not easily result in the accumulation of *X*, because of PPC reaction. If *x* is high, but *y* is also high so that the system is still in the dormant region, PYK plays the same role as the uptake reaction in the previous case. Thus, a futile cycle converting *A* and *B* into *C* is formed in both cases. If the conversion from *A* and *B* into *C* is slow enough, the other reaction, ADK1 (*A* + *C* ⇌ 2*B*) proceeds to balance *A* + *B* and *C*. However, ADK1 cannot balance them if the conversion is too fast because the reaction needs *A*.

Indeed, this intuitive description is consistent with the analytical estimate of the boundary. We derive the boundary between the growth and dormant region for *x* ≪ 1. The boundary is given by *y* leading to a low *a* and *b* with a given value of *x*. For the estimation of the critical *y*, we sum up d*a*/d*t* and d*b*/d*t* and assume *a,b,x* = *O*(*ϵ*) with *ϵ* ≪ 1. Also, recall that the irreversibility parameter *κ* in (Eq. (2)) is small, and thus, we omit *κ* term^2^. Then we obtain

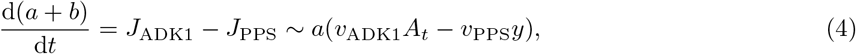

where we explicitly write down the rate parameter *v*\*’s for the interpretation of the estimate. If the first term of the rightmost side is larger than the second term, *a* + *b* increases, while in the opposite situation, the sum keeps decreasing to zero as long as *a* is non-zero. This shift occurs at *v*_PPS_*y* ~ *v*_ADK1_*A_t_*, and it gives the boundary between the growing and the dormant regions.

Next, we explain how the decrease of *a* and *b* leads to the vector field parallel to the horizontal axis in the dormant region (Fig. 4D). Let us assume that *a* and b are approximately the same and well-represented by a single lumped parameter *ϕ*. Also, we set the irreversibility parameter *κ* to zero for simplicity. Then, the ODE for the 2-variable minimal model (Eq. (2)) is given by

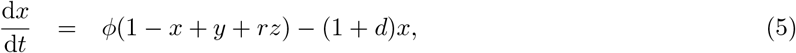

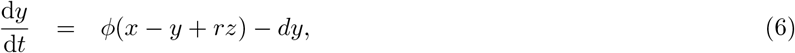

where *z* is the function of *ϕ*, while it becomes constant as *ϕ* approaches 0. From the equation, we can see that if *a* and *b*, represented by *ϕ*, are *O*(1) (i.e., in the growth region), the timescale of the system is *O*(1). On the other hand, if *A* and *B* deplete and *ϕ* ≈ 0 holds in the dormant region, the timescale of d*y*/d*t* becomes *O*(*d*). Since the spontaneous degradation rate *d* is sufficiently smaller than unity, |d*x*/d*t*| ≫ |d*y*/d*t*| holds, and it leads to the vector field being almost parallel to the *x* axis as depicted in Fig. 4D.

To confirm if the simplification above still captures the feature of the vector filed in Fig. 4D, we have drawn the vector field of the simplified model Eq. (5) and (6) with *ϕ* = max{1 – *y, ϕ*_0_} in Fig. 4E. It well captures the feature of the original vector field. We have confirmed that the shape of the vector field is robust to the choice of the function *ϕ*. Also, we analytically solved the model without the growth dilution term (Eq.(5) and (6) with *r* = 0) and found that the model has only a single timescale which is O(1) in the growth region (see *SI Text* Section.4).

The simplified model (Eq.(5) and (6)) highlights that the timescale of dx/dt is much faster than that of d*y*/d*t* in the dormant region. The right hand side of Eq.(5) has the term (1 + *d*)*x*, while that in Eq.(6) is only the degradation term *dy*, and this difference results in the parallel streamline in the phase space (Fig. 4E). It is worth noting where the term (1 + *d*)*x* in Eq.(5) comes from. d corresponds to the constant-rate degradation term, and the reactions coupled with either *A* or *B* should have the rate proportional to *ϕ*. Therefore, this timescale 1 comes from the reaction coupled neither with *A* nor *B*, namely, PPC (*X* → *Z*). All the reactions except PPC are coupled with either *A* or *B*, and thus, the reactions slow down over the boundary between the growth- and the dormant region. However, the rate of PPC has no direct effect from the depletion of *A* and *B*. Then, even after the slowing down of almost all reactions, *X* is kept being consumed, and it leads to the characteristic dynamics of the dormant trajectory.

Note that, if PPC were also coupled with *A* and *B*, (1 + *d*)*x* term in Eq.(5) would have been replaced by (*ϕ* + *d*)*x*. In such a case, all the reactions would have been uniformly slowed down by the depletion of *A* and *B*, and the direction of the vector field would not change over the boundary as drastically as Fig. 4D. Thus, it is vital that the reaction system partially slows down due to the depletion of *A* and *B*.

It is noteworthy that the network structure is also a part of the mechanism: if PPC were the reaction converting *Y* to *Z* instead of *X* to *Z*, the drastic change of the direction of the vector field as Fig. 4D would not result. If PPC were *Y* → *Z*, the main body of the reaction network (reactions except for ADK1) would have no branch. The slowing down of the upstream reactions of PPC (i.e., uptake, PYK, and PPS) would be rate-limiting steps of it, and PPC would slow down coordinated with these reactions.

The above two points suggest that large discrepancies in the chemical concentrations between the steadystate and the plateaux lead to distinct dynamics. In both cases–PPC with energy coupling and the main network without a branch–the reactions uniformly slow down. In such scenarios, even if *A* and *B* deplete, the difference between the production and consumption of each chemical stays relatively small. Thus, the changes in the concentrations remain small. However, if the slowing-down occurs heterogeneously on the network, some chemicals will have a significant mismatch between production and consumption. As a consequence, the concentrations of such chemicals drastically change from the concentrations before the depletion of *A* and *B*.

To sum up, the mechanism of the emergence of the distinct trajectories has two parts: (i) the unbalance of energy (ATP and ADP) production and consumption, and (ii) partial slowing-down of the reaction system caused by non-uniform coupling to the energy currencies and branching of the network.

## Discussion

We have shown that *E. coli* central carbon metabolism exhibits distinctly different dynamics depending on the perturbation from the steady-state concentration. The two types of trajectories greatly differed in terms of the relaxation time and the growth rate during the relaxation, and thus, we termed them as the growth- and the dormant trajectories. We systematically reduce the reaction network without losing the distinct trajectories. By the successive reduction of the model, we eventually reached the minimal network still exhibiting the qualitatively same behavior.

By drawing the vector field of the 2-variable minimal model, we found that there is a boundary at which the vector field changes the direction drastically. Indeed, the two regions are divided by the boundary corresponding to the set of the initial points of the growth- and the dormant trajectories. The analysis led that there are at least two vital requirements for the distinct trajectories: (i) the unbalance of the energy production and consumption and (ii) the partial slowing-down of the reactions due to the non-uniform coupling with the energy currency molecules and branching of the network.

By studying several model variants, we carefully examined the robustness of our main results, namely, the emergence of distinct trajectories and the consequence of model reduction. First, the robustness of the emergence of distinct trajectories to the parameter values was examined. For several values of the total adenine nucleotide carriers concentration (*A_t_*), we randomly assigned the rate constants and studied if the distinct trajectories still emerged. As anticipated from the analysis, the distinct trajectories robustly emerge as long as the total concentration of adenine carriers, *A_t_*, is not too large for the depletion of ATP and ADP to occur (SI *Text* Section.8). As well as the random parameter values, we performed the reaction removal for the model reduction in 16 different randomized orders. All the minimal networks obtained by the random reduction were larger than those derived in the result section but shared the main features argued above, namely, (a) the models keep ADK1 reaction and AMP, and (b) in each network, there are both reactions, with- and without the coupling to the energy currency molecules (ATP, ADP, and AMP) as well as branches.

We checked the outcomes of relaxing the assumptions that we made initially. It is confirmed that the model without the replacement of the nicotinamide nucleotide carriers by the adenine nucleotide carriers exhibits distinct trajectories (SI *text* Section.6). Also, we have confirmed that the distinct trajectories emerge if the assumption on the constant total concentration of the adenine nucleotide carriers is relaxed by introducing a phenomenological reaction for the de-novo synthesis of AMP to the minimal model (SI *Text* Section.5). Overall, the emergence of distinct trajectories is a robust feature of the *E. coli* core model rather than a phenomenon led by fine-tuning of the parameters.

Two transition paths to the dormancy were schematically proposed by M. Heinemann’s group [11,13]. The first path is the transition triggered by stochastic gene expression. For example, the elevated concentration of the toxin protein inhibits several cellular processes and leads to the ppGpp-mediated stress responses [41,42].

There are multiple mathematical models in accordance with this “genetically-triggered” scenario [14,43–47]. The second path is “metabolically-triggered”. The disturbance of the metabolic state induces the stress response to modulate the gene expression pattern. While the disturbance of the metabolic state is attributed to the stochastic fluctuations of the enzyme level in the papers [11, 13], the distinct dynamics due to the nonlinearity of the metabolic reactions should be able to play a role in the transition.

The present model showed a possibility that the dormancy transition could be triggered by the metabolic dynamics itself when the metabolic state is perturbed. According to the minimal model, the perturbation evokes a futile cycle and leads to the depletion of ATP and ADP. Here, the sources of the perturbation can be starvation, nutrient shift, exposure to antibiotics, pH stress, or even stochasticity of the intracellular processes.

However, in reality, the intracellular states of the dormant cells may go beyond what we can depict in terms of metabolites. We did not consider a dynamic change in enzyme concentrations in the current model analysis, which would modify chemical reaction rates. While a constant enzyme concentration assumption is plausible in steady-state growth, gene regulations of enzyme levels are likely to be relevant in dynamically changing growth processes. Indeed, disruptions of the metabolic states may lead to several responses, such as the stress-response systems controlled by (p)ppGpp, toxin-antitoxin modules, and/or the alternative sigma factor *σ^S^* [11,48–53]. Since the response systems work to relieve the disruption of the metabolic state, such a strong drop of the concentrations of the metabolites found in the present study may be avoided by gene regulation of relevant enzymes. While further research on the effect of the gene-regulation dynamics is needed, we believe that the present findings provide a possible mechanism to trigger dormancy where the disruption of the metabolic state may lead to even bigger responses.

Note that we can find the counterpart of the reactions in the minimal model in the full *E.coli* metabolic network. Thus, the dormancy transition demonstrated by the minimal model is verifiable by experiments. The central part of the mechanism is that PPS and PYK can form a futile cycle and the competition between PPS and ADK1 on the consumption/production of ATP. Indeed, the experiments showed that one could induce the ATP-consuming futile cycle between phosphoenolpyruvate and pyruvate via PYK and PPS by overexpressing the ppsA gene [54, 55]. Taking the experimental reports and the present computational results together, we hypothesize that the overexpression of the pps gene leads to an increase in the persister fraction because PPS converts ATP to AMP.

Lastly, we like to remark that the requirements for the metabolic network to show the distinct trajectories are not limited only on the *E. coli* core model. We can find several reactions that potentially form a futile cycle from various species. For instance, each of Acetyl-CoA synthetase (KEGGID:R00235), Phosphoribosylpyrophosphate synthetase (R01049), and Asparagine synthase (R00578) [56] converts ATP to AMP and forms a loop in the metabolic networks. These are the minimum requirements for a reaction to form a futile cycle discussed above. Such reactions are widespread from prokaryotes to eukaryotes and from unicellular to multicellular organisms. Comprehensive studies of the kinetic models of not only *E. coli* but also other organisms may pave the way for understanding the robust and generic network features leading to the multiple timescales of cellular growth and dormancy.

## Acknowledgments

The authors thank Chikara Furusawa and Sandeep Krishna for the fruitful discussion. This work is supported by the research grants from VILLUM FONDEN (00028054) and the Novo Nordisk Foundation (NNF21OC0068775).

## Materials and Methods

### Simulation of ordinary differential equations (ODEs)

All the ODE computations were performed by using Matlab (Mathworks) `ode23s` function. For searching attractors, we set 10^*u*_*i,n*_^ for the *i*th metabolite as the nth initial value where *u_i,n_* is the random number generated from a uniform distribution in [–1,1]. The steady-state concentration 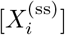 is then obtained, initial concentrations for the main analysis of the dynamics are generated as 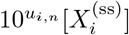 with *u_i,n_* is the same random number yet distributed in [–2, 2]. The ODEs were computed with two tolerance options (`AbsTol` = 10^−10^, `RelTol` = 10^−12^) and (`AbsTol` = 10^−10^, `RelTol` = 10^−14^) from exactly the same initial points. After the computation, the trajectories with two different `RelTol` values, but from the same initial point, were compared for the quality check of the computation. If the Hausdorff distance of the pair of the trajectories was less than 0.5, the trajectories were considered as correctly computed, and the trajectory obtained with `RelTol` = 10^−14^ was used for the further analysis, and otherwise, discarded. The quality check of the computation was performed after the transformation *x*(*t*) → ln(*x*(*t*)) where *x*(*t*) is the concentration of the chemicals.

### Principal Component Analysis

We used the python package `sklearn.decomposition.PCA` [57] without whitening. The whitening leads to only a minor effect on the results. The concentrations of the chemicals were transformed into the natural logarithm of the concentration before the analysis. PCA is performed for each model, i.e., all the data points generated by a single model with a number of initial conditions (the state vectors representing the chemical concentrations) are stacked into a single dataset, and we computed the covariance matrix of the dataset for the projection of the trajectories.

## Footnotes

1 The concentration of glutamine (the purple bottommost line in the top panel of A) reached unrealistically low value even in the steady-state. We attribute this behavior to the following technical reason: The biomass synthesis reaction was not incorporated into the model in [33] where the kinetic parameters were estimated. In the present model, we added the biomass synthesis reaction to consider the cell growth, which resulted in much faster consumption of glutamine than in the model used for parameter estimate. See *SI Text* Section1 for details.

2 This is not necessary for the argument, but makes the equation complicated.

## References

[1] Guennadi Sezonov, Danièle Joseleau-Petit, and Richard d’Ari. Escherichia coli physiology in luria-bertani broth. Journal of bacteriology, 189(23):8746–8749, 2007.

[2] Nathalie Q Balaban, Jack Merrin, Remy Chait, Lukasz Kowalik, and Stanislas Leibler. Bacterial persistence as a phenotypic switch. Science, 305(5690):1622–1625, 2004.

[3] Nathalie Q Balaban, Sophie Helaine, Kim Lewis, Martin Ackermann, Bree Aldridge, Dan I Andersson, Mark P Brynildsen, Dirk Bumann, Andrew Camilli, James J Collins, et al. Definitions and guidelines for research on antibiotic persistence. Nature Reviews Microbiology, 17(7):441–448, 2019.

[4] Susanne K Christensen, Marie Mikkelsen, Kim Pedersen, and Kenn Gerdes. Rele, a global inhibitor of translation, is activated during nutritional stress. Proceedings of the National Academy of Sciences, 98(25):14328–14333, 2001.

[5] Yusuke Himeoka and Namiko Mitarai. When to wake up? the optimal waking-up strategies for starvation-induced persistence. PLoS computational biology, 17(2):e1008655, 2021.

[6] Mikkel Skjoldan Svenningsen, Sine Lo Svenningsen, Michael Askvad Sørensen, and Namiko Mitarai. Existence of log-phase escherichia coli persisters and lasting memory of a starvation pulse. Life Science Alliance, 5(2), 2022.

[7] Irit Levin-Reisman, Orit Gefen, Ofer Fridman, Irine Ronin, David Shwa, Hila Sheftel, and Nathalie Q Balaban. Automated imaging with scanlag reveals previously undetectable bacterial growth phenotypes. Nature Methods, 7(9):737, 2010.

[8] Yusuke Himeoka and Kunihiko Kaneko. Theory for transitions between exponential and stationary phases: universal laws for lag time. Physical Review X, 7(2):021049, 2017.

[9] Yusuke Himeoka, Bertil Gummesson, Michael A Sørensen, Sine Lo Svenningsen, and Namiko Mitarai. Distinct survival, growth lag, and rRNA degradation kinetics during Long-Term starvation for carbon or phosphate. mSphere, page e0100621, April 2022.

[10] Elena Biselli, Severin Josef Schink, and Ulrich Gerland. Slower growth of escherichia coli leads to longer survival in carbon starvation due to a decrease in the maintenance rate. Molecular Systems Biology, 16(6):e9478, 2020.

[11] Jakub Leszek Radzikowski, Silke Vedelaar, David Siegel, Álvaro Dario Ortega, Alexander Schmidt, and Matthias Heinemann. Bacterial persistence is an active σs stress response to metabolic flux limitation. Molecular systems biology, 12(9):882, 2016.

[12] Stephanie M Amato, Christopher H Fazen, Theresa C Henry, Wendy W K Mok, Mehmet A Orman, Elizabeth L Sandvik, Katherine G Volzing, and Mark P Brynildsen. The role of metabolism in bacterial persistence. Front. Microbiol., 5:70, March 2014.

[13] Jakub Leszek Radzikowski, Hannah Schramke, and Matthias Heinemann. Bacterial persistence from a system-level perspective. Current Opinion in Biotechnology, 46:98–105, 2017.

[14] Stefan Klumpp, Zhongge Zhang, and Terence Hwa. Growth rate-dependent global effects on gene expression in bacteria. Cell, 139(7):1366–1375, 2009.

[15] Ido Golding, Johan Paulsson, Scott M Zawilski, and Edward C Cox. Real-time kinetics of gene activity in individual bacteria. Cell, 123(6):1025–1036, December 2005.

[16] Keisuke Fujita, Mitsuhiro Iwaki, and Toshio Yanagida. Transcriptional bursting is intrinsically caused by interplay between RNA polymerases on DNA. Nat. Commun., 7:13788, December 2016.

[17] Akinori Awazu and Kunihiko Kaneko. Ubiquitous “glassy” relaxation in catalytic reaction networks. Physical Review E, 80(4):041931, 2009.

[18] Akinori Awazu and Kunihiko Kaneko. Discreteness-induced slow relaxation in reversible catalytic reaction networks. Physical Review E, 81(5):051920, 2010.

[19] Tetsuhiro S Hatakeyama and Chikara Furusawa. Metabolic dynamics restricted by conserved carriers: Jamming and feedback. PLoS computational biology, 13(11):e1005847, 2017.

[20] Christophe Chassagnole, Naruemol Noisommit-Rizzi, Joachim W Schmid, Klaus Mauch, and Matthias Reuss. Dynamic modeling of the central carbon metabolism of escherichia coli. Biotechnology and bioengineering, 79(1):53–73, 2002.

[21] Hiroyuki Kurata and Yurie Sugimoto. Improved kinetic model of escherichia coli central carbon metabolism in batch and continuous cultures. Journal of bioscience and bioengineering, 125(2):251–257, 2018.

[22] Tuty Asmawaty Abdul Kadir, Ahmad A Mannan, Andrzej M Kierzek, Johnjoe McFadden, and Kazuyuki Shimizu. Modeling and simulation of the main metabolism in escherichia coli and its several single-gene knockout mutants with experimental verification. Microbial cell factories, 9(1):1–21, 2010.

[23] Yukako Tohsato, Kunihiko Ikuta, Akitaka Shionoya, Yusaku Mazaki, and Masahiro Ito. Parameter optimization and sensitivity analysis for large kinetic models using a real-coded genetic algorithm. Gene, 518(1):84–90, 2013.

[24] Oliver Kotte, Judith B Zaugg, and Matthias Heinemann. Bacterial adaptation through distributed sensing of metabolic fluxes. Molecular systems biology, 6(1):355, 2010.

[25] Kirill Peskov, Ekaterina Mogilevskaya, and Oleg Demin. Kinetic modelling of central carbon metabolism in escherichia coli. The FEBS journal, 279(18):3374–3385, 2012.

[26] Ahmad A Mannan, Yoshihiro Toya, Kazuyuki Shimizu, Johnjoe McFadden, Andrzej M Kierzek, and Andrea Rocco. Integrating kinetic model of e. coli with genome scale metabolic fluxes overcomes its open system problem and reveals bistability in central metabolism. PloS one, 10(10):e0139507, 2015.

[27] Jimena Di Maggio, JC Diaz Ricci, and M Soledad Diaz. Global sensitivity analysis in dynamic metabolic networks. Computers & Chemical Engineering, 34(5):770–781, 2010.

[28] Katja Bettenbrock, Sophia Fischer, Andreas Kremling, Knut Jahreis, Thomas Sauter, and Ernst-Dieter Gilles. A quantitative approach to catabolite repression in escherichia coli. Journal of Biological Chemistry, 281(5):2578–2584, 2006.

[29] Vivek Kumar Singh and Indira Ghosh. Kinetic modeling of tricarboxylic acid cycle and glyoxylate bypass in mycobacterium tuberculosis, and its application to assessment of drug targets. Theoretical Biology and Medical Modelling, 3(1):1–11, 2006.

[30] C Juan Torres, Victoria Guixé, and Jorge Babul. A mutant phosphofructokinase produces a futile cycle during gluconeogenesis in escherichia coli. Biochemical Journal, 327(3):675–684, 1997.

[31] Yikun Tan and James C Liao. Metabolic ensemble modeling for strain engineers. Biotechnology journal, 7(3):343–353, 2012.

[32] Linh M Tran, Matthew L Rizk, and James C Liao. Ensemble modeling of metabolic networks. Biophysical journal, 95(12):5606–5617, 2008.

[33] Ali Khodayari, Ali R Zomorrodi, James C Liao, and Costas D Maranas. A kinetic model of escherichia coli core metabolism satisfying multiple sets of mutant flux data. Metabolic engineering, 25:50–62, 2014.

[34] Ali Khodayari and Costas D Maranas. A genome-scale escherichia coli kinetic metabolic model k-ecoli457 satisfying flux data for multiple mutant strains. Nature communications, 7(1):1–12, 2016.

[35] Matthew L Rizk and James C Liao. Ensemble modeling for aromatic production in escherichia coli. PloS one, 4(9):e6903, 2009.

[36] Sylvie Manuse, Yue Shan, Silvia J Canas-Duarte, Somenath Bakshi, Wei-Sheng Sun, Hirotada Mori, Johan Paulsson, and Kim Lewis. Bacterial persisters are a stochastically formed subpopulation of low-energy cells. PLoS biology, 19(4):e3001194, 2021.

[37] Brian P Conlon, Sarah E Rowe, Autumn Brown Gandt, Austin S Nuxoll, Niles P Donegan, Eliza A Zalis, Geremy Clair, Joshua N Adkins, Ambrose L Cheung, and Kim Lewis. Persister formation in staphylococcus aureus is associated with atp depletion. Nature microbiology, 1(5):1–7, 2016.

[38] Yue Shan, Autumn Brown Gandt, Sarah E Rowe, Julia P Deisinger, Brian P Conlon, and Kim Lewis. Atp-dependent persister formation in escherichia coli. MBio, 8(1), 2017.

[39] Palsson B Orth J, Fleming R. Reconstruction and Use of Microbial Metabolic Networks: the Core Escherichia coli Metabolic Model as an Educational Guide. 2010.

[40] Zachary A King, Andreas Dräger, Ali Ebrahim, Nikolaus Sonnenschein, Nathan E Lewis, and Bernhard O Palsson. Escher: a web application for building, sharing, and embedding data-rich visualizations of biological pathways. PLoS Comput Biol, 11(8):e1004321, 2015.

[41] Devang Shah, Zhigang Zhang, Arkady Khodursky, Niilo Kaldalu, Kristi Kurg, and Kim Lewis. Persisters: a distinct physiological state of e. coli. BMC Microbiol., 6: 53, June 2006.

[42] Shaleen B Korch and Thomas M Hill. Ectopic overexpression of wild-type and mutant hipa genes in escherichia coli: effects on macromolecular synthesis and persister formation. J. Bacteriol., 188(11):3826–3836, June 2006.

[43] Eitan Rotem, Adiel Loinger, Irine Ronin, Irit Levin-Reisman, Chana Gabay, Noam Shoresh, Ofer Biham, and Nathalie Q Balaban. Regulation of phenotypic variability by a threshold-based mechanism underlies bacterial persistence. Proceedings of the National Academy of Sciences, 107(28):12541–12546, 2010.

[44] Andrea Rocco, Andrzej M Kierzek, and Johnjoe McFadden. Slow protein fluctuations explain the emergence of growth phenotypes and persistence in clonal bacterial populations. PloS one, 8(1):e54272, 2013.

[45] Ilaria Cataudella, Kim Sneppen, Kenn Gerdes, and Namiko Mitarai. Conditional cooperativity of toxin-antitoxin regulation can mediate bistability between growth and dormancy. PLoS computational biology, 9(8):e1003174, 2013.

[46] Lendert Gelens, Lydia Hill, Alexandra Vandervelde, Jan Danckaert, and Remy Loris. A general model for toxin-antitoxin module dynamics can explain persister cell formation in e. coli. PLoS computational biology, 9(8):e1003190, 2013.

[47] Jingchen Feng, David A Kessler, Eshel Ben-Jacob, and Herbert Levine. Growth feedback as a basis for persister bistability. Proceedings of the National Academy of Sciences, 111(1):544–549, 2014.

[48] Dao Nguyen, Amruta Joshi-Datar, Francois Lepine, Elizabeth Bauerle, Oyebode Olakanmi, Karlyn Beer, Geoffrey McKay, Richard Siehnel, James Schafhauser, Yun Wang, Bradley E Britigan, and Pradeep K Singh. Active starvation responses mediate antibiotic tolerance in biofilms and nutrient-limited bacteria. Science, 334(6058):982–986, November 2011.

[49] Vasili Hauryliuk, Gemma C Atkinson, Katsuhiko S Murakami, Tanel Tenson, and Kenn Gerdes. Recent functional insights into the role of (p)ppgpp in bacterial physiology. Nat. Rev. Microbiol., 13(5):298–309, May 2015.

[50] Regine Hengge-Aronis. Signal transduction and regulatory mechanisms involved in control of the σs (rpos) subunit of rna polymerase. Microbiology and molecular biology reviews, 66(3):373–395, 2002.

[51] Rv Lange and Regine Hengge-Aronis. Identification of a central regulator of stationary-phase gene expression in escherichia coli. Molecular microbiology, 5(1):49–59, 1991.

[52] Rebecca Page and Wolfgang Peti. Toxin-antitoxin systems in bacterial growth arrest and persistence. Nature chemical biology, 12(4):208–214, 2016.

[53] Pooja Sharma, Parth Pratim Pandey, and Sanjay Jain. Modeling the cost and benefit of proteome regulation in a growing bacterial cell. Phys. Biol., 15(4):046005, May 2018.

[54] R Patnaik, WD Roof, RF Young, and JC Liao. Stimulation of glucose catabolism in escherichia coli by a potential futile cycle. Journal of bacteriology, 174(23):7527–7532, 1992.

[55] Oliver Hädicke, Katja Bettenbrock, and Steffen Klamt. Enforced atp futile cycling increases specific productivity and yield of anaerobic lactate production in escherichia coli. Biotechnology and bioengineering, 112(10):2195–2199, 2015.

[56] Minoru Kanehisa and Susumu Goto. Kegg: kyoto encyclopedia of genes and genomes. Nucleic acids research, 28(1):27–30, 2000.

[57] F. Pedregosa, G. Varoquaux, A. Gramfort, V. Michel, B. Thirion, O. Grisel, M. Blondel, P. Prettenhofer, R. Weiss, V. Dubourg, J. Vanderplas, A. Passos, D. Cournapeau, M. Brucher, M. Perrot, and E. Duchesnay. Scikit-learn: Machine learning in Python. Journal of Machine Learning Research, 12:2825–2830, 2011.

