## Supplementary material for "Emergence of growth and dormancy from a kinetic model of the *Escherichia coli* central carbon metabolism": SI Text

**S4 Analytic solution of the simple model and the choice of the function $\phi$** **12**

**S5 The minimal model with de-novo AMP synthesis** **13**

**S6 model0 with the nicotinamide nucleotide carriers** **15**

**S7 Model reduction with random order** **16**

**S8 The behavior of model0 with randomly assigned parameters** **19**

#### S1 The *E. coli* core model with biologically realistic parameter values

It is worth asking if the model exhibits distinct trajectories with biologically realistic kinetic values. However, obtaining the kinetic parameters for all reactions, even in a rather small *E. coli* core model, is still challenging. Thus, we take advantage of the metabolic ensemble modeling (MEM) [1, 2], which is a method for the parameter estimation of the metabolic models. In the MEM approach, each enzymatic reaction is decomposed into a sequence of elementary reactions, i.e., an enzymatic reaction $A+B \rightleftharpoons C$ catalyzed by $E$ is, for instance, decomposed as follows

$$A + E \rightleftharpoons AE \quad \text{(S1)}$$
$$AE + B \rightleftharpoons ABE \quad \text{(S2)}$$
$$ABE \rightleftharpoons CE \quad \text{(S3)}$$
$$CE \rightleftharpoons C + E \quad \text{(S4)}$$

Note that the rates of elementary reactions (Eq.(S1)-(S4)) can be represented by the mass-action kinetics. For example, the forward reaction of Eq.(S1) is given by

$$v_{A+E\to AE} = k_{A+E\to AE}[A][E],$$

where $k_{A+E\to AE}$, $[A]$, and $[E]$ represents the rate-constant of the reaction $A + E \to AE$, the concentration of the chemical $A$, and the concentration of the free enzyme $E$ ($E$ not in the complex form). Let $[A]_{ss}$ and $[E]_0$ be the steady-state concentration of the chemical $A$ and the total concentration of the enzyme $E$, respectively. Then at the steady state, the logarithm of the reaction flux is

$$\ln v_{A+E\to AE} = \ln(k_{A+E\to AE}[A]_{ss}[E]_0) + \ln([E]/[E]_0).$$

Note that the term depending on $[A]$ is dropped because $\ln[A]/[A]_{ss}$ is zero at the steady-state. The MEM approach seeks the values of the scaled rate-constant such as $\tilde{k}_{A+E\to AE} = k_{A+E\to AE}[A]_{ss}[E]_0$ and $e = [E]/[E]_0$ so that the model can fit the experimentally-obtained fluxome data using the ensemble modeling [3]. (for more detail, see [1,2])

For the simulation of the *E. coli* core model, we adopted the parameter values estimated by Khodayari et al. [2]. For obtaining the values of (non-scaled) rate constants, we need to divide the scaled rate constants by experimentally reported concentrations of chemicals because what they estimated are, for instance, in the form of $k_{A+E\to AE}[A]_{ss}[E]_0$. We calculated the rate constants by using the concentration data measured by Gerosa et al. [4] and estimated by Akbari et al. [5]. Since the concentration of glyoxylate was presented in neither [4] nor [5], we used the geometric mean of the concentrations of two neighbor metabolites in the metabolic network, isocitrate and L-malate. The back-calculated parameters are presented in *SI Data.*2. After the back-calculation of the rate constants, we constructed an ODE model. We adiabatically eliminated the elementary reactions for each enzymatic reaction in the model and used the Michaelis-Menten type rate equation (see [2]).

In growth dynamics (Fig. 2A in the main text), the concentration of a chemical species, glutamine, becomes lower than 1nM. Glutamine is one of the growth factors in the current setup. We attribute this extreme drop to the following technical reason: In contrast to the present model, the biomass synthesis reaction was not incorporated into the model in [2] where the kinetic parameters were estimated. Therefore, glutamine is consumed much faster in the present model than in the model used to estimate the parameters.

### S2 Simplified model with mass-action kinetics

The kinetic *E. coli* core model is, as it is, too complicated to understand the mechanism that leads to the two distinct relaxation trajectories. Thus, we simplified the *E. coli* core model as follows. First, we modified the kinetics of the chemical reactions from the Michaelis-Menten formula to the mass-action rate equation. The rate of the $i$th chemical reaction $A \rightleftharpoons B$, $J_i$, which was given by

$$J_i = v_i \frac{[A] - k_i[B]}{1 + [A]/K_A^{(i)} + [B]/K_B^{(i)}} \tag{S5}$$

is replaced by

$$J_i = v_i([A] - k_i[B]), \tag{S6}$$

where $v_i$ and $v_i k_i$ are the forward- and backward reaction rate constant, respectively. Note that the mass-action kinetics (Eq.(S6)) is a special form of the Michaelis-Menten kinetics (Eq.(S5)) in the parameter region where $[A] \ll K_A^{(i)}$ and $[B] \ll K_B^{(i)}$ hold (for general arguments, see [6]). The model is then nondimensionalized by scaling the concentrations by the external glucose concentration and the time by the rate constant of the glucose uptake. We further simplified the rate equations by setting $v_i$'s to unity and binarising $k_i$'s for all $i'$s. The *E. coli* core model contains the information of irreversibility for each reaction, and thus, if the $i$th reaction is reversible, we set $k_i$ as unity, and otherwise, set it to $\kappa \ll 1$. We term this simplified version of the kinetic *E. coli* core model as model0 with an index for the following model reduction steps.

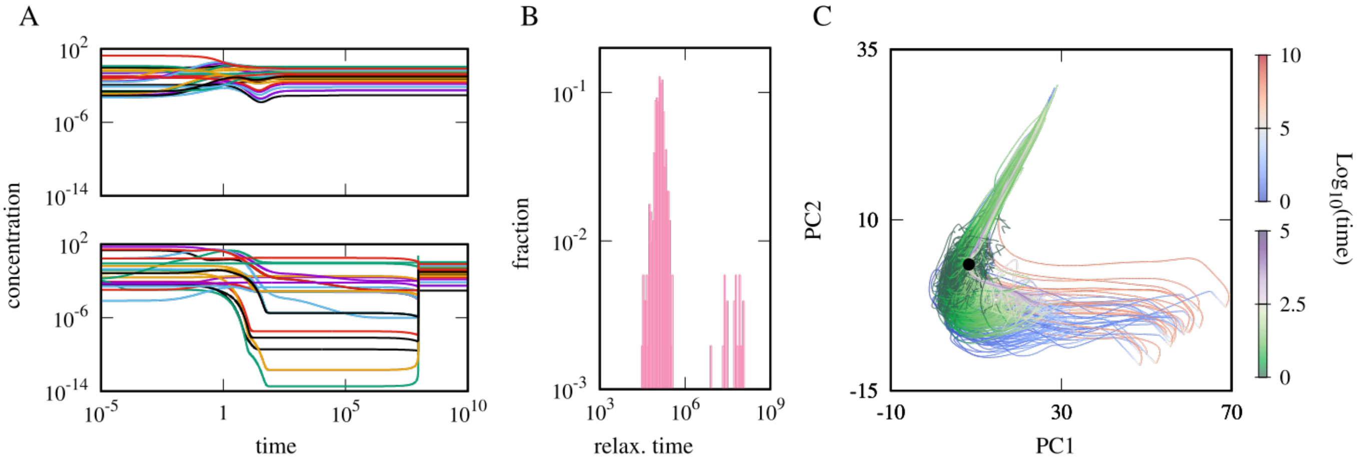

Figure S1: A. Two characteristic dynamics of the model0 starting from different initial points. The relaxation behaviors are qualitatively different between the top and bottom panels. B. The distribution of the relaxation time showing a clear bimodality. C. Trajectories are overlaid in 2-dimensional principal component space. The color indicates $\log_{10}$ of time. The trajectories having shorter relaxation time (top panel of A) are colored green-white-purple while the others are colored blue-white-red. The black point corresponds to the attractor. Initial concentration of each metabolites is $10^{u_{i,n}}[X_i^{(\mathrm{ss})}]$ with $[X_i^{(\mathrm{ss})}]$ as the steady-state concentration of the $i$th metabolite, and $u_{i,n}$ as a random number uniformly distributed in $[-2, 2]$ while the total concentrations of adenine nucleotide carriers are normalized. $v = 1$ and $\kappa = 10^{-6}$ for all reactions. Other parameters are $[\mathrm{glc}] = 1, A_t = 1, r = 0.1$ and $d = 10^{-8}$.

Surprisingly, the emergence of distinct relaxation trajectories is robust to such an extensive model modification. The qualitative difference of the trajectories (Figs. S1A), bimodality of the distribution of the relaxation time (Fig. S1B), and the distinction of the trajectories in the PC1-PC2 space (Fig. S1C) were unchanged. This robustness implies that the emergence of the distinct trajectories stems from the structure of the metabolic reaction network of the *E. coli* core model rather than choices of specific parameter values. We also confirmed that the distinct trajectories emerge if the kinetic parameters are randomly assigned instead of setting them to unity (see Section S8).

### S3 Model Reduction

#### S3.1 Criterion for distinct trajectories

For each model reduction step, we run the ODE model from 512 randomly generated initial points to search the attractors. Except for a single model, all the intermediate models of the reduction process showed monostability (see SectionS3.6). Then, 512 initial conditions were generated by perturbing the steady-state concentration in the same way as we did in the previous models.

The bimodality of the relaxation time distribution is one of the best intuitive criteria for distinct trajectories. However, we found in some cases, the bimodality was unclear even though there were clearly different types of trajectories when we plotted the time courses and performed PCA. This is because the growth rates during the relaxation of both types of trajectories (growth and dormant) become smaller than the spontaneous degradation rate $d$. Thus, the relaxation time of all the trajectories becomes approximately $1/d$. It is possible to overcome this problem by setting $d$ to be sufficiently small such as $10^{-20}$ in principle, but the computation time becomes unbearably long. Thus, we decided to focus on the similarity of the trajectories instead of the relaxation time itself. For interested readers, the distributions of the relaxation time for all the models in the reduction step are presented in Section S3.3.

Below, we first intuitively explain how we quantify the similarity of the trajectories and then introduce the actual measure.

Suppose that a model has a single attractor. Then, all the trajectories starting from different initial points eventually converge. We like to categorize the trajectories into different groups so that if a pair of trajectories monotonically approach each other as they converge to the attractor, they belong to the same group. One may naïvely expect that we can state that two trajectories $x(t)$ and $y(t)$ monotonically approach each other if the Euclidean distance between them at the same time point, $d(x(t), y(t))$, is a monotonically decreasing function of $t$. However, since the initial points are distributed in the phase space, measuring the distance between the points on two trajectories at the same time point is unreasonable.

Thus, instead of adopting this naïve definition of monotonicity, we measure the maximum Euclidean distance between two trajectories in the phase space. It is known as the Hausdorff distance of the trajectories, given by

$$d_H(x, y) = \max\{\max_t \min_s d(x(t), y(s)), \max_t \min_s d(x(s), y(t))\}. \tag{S7}$$

The Hausdorff distance first looks for the closest point of the trajectory $y$ from the point $x(t)$, $y(s^*(t))$, and then find the pair of the points $(x(t), y(s^*(t)))$ which gives the maximum Euclidean distance. The same is done from the points of $y(t)$ and the larger value is chosen for the symmetry $d_H(x, y) = d_H(y, x)$. The Hausdorff distance thus measures how far the two trajectories are distant while trivially distant pairs of points are not taken into account (for example, the initial point of $x$ and the endpoint of $y$, i.e., the attractor).

We cannot judge whether the trajectories go away from each other or not directly from the Hausdorff distance since it needs to be compared with the initial separation. Thus, we normalize the Hausdorff distance by the Euclidean distance between the initial points, $d(x(0), y(0))$ leading to

$$R(x, y) = \frac{d_H(x, y)}{d(x(0), y(0))}. \tag{S8}$$

We call this ratio $R(x, y)$ as the expansion ratio of the trajectories $x$ and $y$. It measures how much the initial distance has expanded. If $d_H(x, y)$ is smaller than the initial distance, $R(x, y)$ is less than unity. $R(x, y) > 1$ means that two trajectories go away from each other at least once despite eventually converging to the same attractor. Note that in this manuscript, the Euclidean distance, and accordingly, the Hausdorff distance are measured in the original high-dimensional phase space after applying the logarithm-conversion of the variables, not in the lower-dimensional principal component space.

The distribution is expected to have a trivial peak around $R = 1$. If the distribution has only a trivial peak, it indicates that all the trajectories are monotonically attracted to a single predominant stream in the phase space reaching the attractor. Oppositely, suppose the distribution has a non-trivial peak(s) and/or an additional long tail. In that case, the correlation between the initial distance and the Hausdorff distance is not simply scaled to each other.

Therefore, in the present manuscript, we utilize the multimodality and/or the long tail of the distribution of the expansion ratio as the criterion of the distinct trajectories (examples can be found in Section S3.3). We examine if the model has exhibited distinct trajectories by fitting the distribution by a sum of Gaussian functions. In rough terms, it checks if the distribution needs at least two Gaussian functions with distant peaks (see SectionS3.2 for the details). Note that in the following analysis, the computation of the expansion ratio and PCA were performed for the trajectories converted to the logarithmic scale so that the dynamic behaviours of the chemicals with low concentrations are also reflected in the analysis (As a side effect of the logarithm-conversion, the behaviours of the chemicals with quite low concentrations may be too much highlighted. We computed the expansion ratio with cut-offs of the concentrations for a lower side, see SectionS3.4).

#### S3.2 Judging multimodality

Let us suppose that there is a list of the expansion ratio $\{R(x, y)\}_{x,y \in \mathcal{T}_i}$ where $\mathcal{T}_i$ is the set of the trajectories of the $i$th model. Then, we fit the histogram of the expansion ratio by a sum of the Normal distributions

$\mathcal{G}(R, \vec{\mu}, \vec{\sigma}) = \sum_{i=0}^{M-1} w_i \mathcal{N}(R; \mu_i, \sigma_i), \sum_{i=0}^{M-1} w_i = 1, w_i \geq 0$ where $\mathcal{N}(R; \mu_i, \sigma_i)$ is the Normal distribution with $\mu_i$ and $\sigma_i$ as the mean and the standard deviation, respectively. Here, we heuristically choose $M$ as 4 because the distributions of the expansion ratio often had a heavy tail, and fitting with a small $M$ could prioritize to fit the tail rather than the second peak. We used the python package `sklearn.mixture for the fitting.GaussianMixture` [7] with the options as `covariance_type` $=$ `'full'`, `tol` $= 10^{-4}$, `n_init` $= 16$.

After the fitting, we reorganize the indices of the normal distributions so that $\mu_i < \mu_{i+1}$ holds. We judged the distribution is multimodal if the result fulfills the conditions below

- $\mu_0 < 1.05$ (there is a trivial peak)
- $w_0 \geq w_i$ (the trivial peak has the largest weight)
- $1 \leq {}^{\exists}i < M$ s.t., $\mu_i - \mu_0 > \max(1, \sigma_0 + \sigma_i)$ and $w_i/w_0 > 0.01$ (there is another, distant peak)

#### S3.3 Intermediate models

Here, we present the trajectories on PC1-PC2 space (Figs. S2 and S3), the distribution of the expansion ratio (Figs. S4), and the relaxation time distribution (Figs. S5) for all the intermediate models of the reduction described in the main text. As mentioned in the main text, $d = 10^{-8}$ becomes larger than the growth rate $\mu$ at the relaxation plateaux for some models, and in such cases, the relaxation time cannot distinguish the growth- and dormant trajectories. According to the importance of [atp] + [adp] that we found in the main text, we wonder if the minimum value of [atp] + [adp] during the relaxation of each trajectory works as a criterion to distinguish the two types of the trajectories. In the accordance with the expectation, we found that the distribution of $A_{\min} = \min_{t\in(0,\infty)} \log_{10}([\text{atp}](t) + [\text{adp}](t))$ of each intermediate model was double-peaked. Thus, we colored each trajectory in Figs. S2 and S3 based on which peak of the distribution $A_{\min}$ the trajectory belongs to.

#### S3.4 The expansion ratio with cutoffs

As shown in Figs.2A and Figs.3A, the concentrations of some chemicals become too low. Because of the logarithm conversion of the concentrations, these low concentrations can strongly contribute to the multimodal distributions of the expansion ratio to result. To check if the multimodal distribution of the expansion is sensitive to such low concentrations, we computed the expansion ratio of the model0 with cut-offs. With a given value of cutoff, $C$, we converted each element of the trajectories $\vec{x}(t)$ to $\xi_i(t) = \max(x_i(t), C)$. The trajectories $\vec{\xi}(t)'$s are logarithm-converted and then used for computing the expansion ratio. As shown in Fig.S6, the distributions are multimodal up to $C = 10^{-10}$, while the distribution becomes long-tailed with two plateaus for $C \gtrapprox 10^{-9}$.

#### S3.5 $L/D$ ratio

The trajectories projected onto the two-dimensional PC space give us the impression that the dormant trajectories take roundabout ways comparing the growth trajectories. For the confirmation of the impression, we compare the length of the trajectory in the phase space.

For a trajectory $x(t)$, we introduce two quantities, namely, the line integral of the trajectory $L = \int_x dl$, and the Euclidean distance between the initial point and the attractor $D = d(x(0), x(\infty))$. Since the straight line gives the shortest possible length between two points, the ratio $L/D$ of $x$ measures the deviation of the trajectory $x$ from the shortest path from the initial point to the attractor, representing how far $x$ takes a detour.

For grouping the trajectories, we used the minimum value of [atp] + [adp] during the relaxation $A_{\min}$ of each trajectory (see SectionS3.3). We computed the average $L/D$ ratio of the high $A_{\min}$ (growth) and the low $A_{\min}$ (dormant) trajectories, respectively. As shown in Fig. S7, the average $L/D$ ratio of the low $A_{\min}$ trajectories is larger than that of the trajectories with high $A_{\min}$ values for all the models while the differences are within the error bar in model 5.

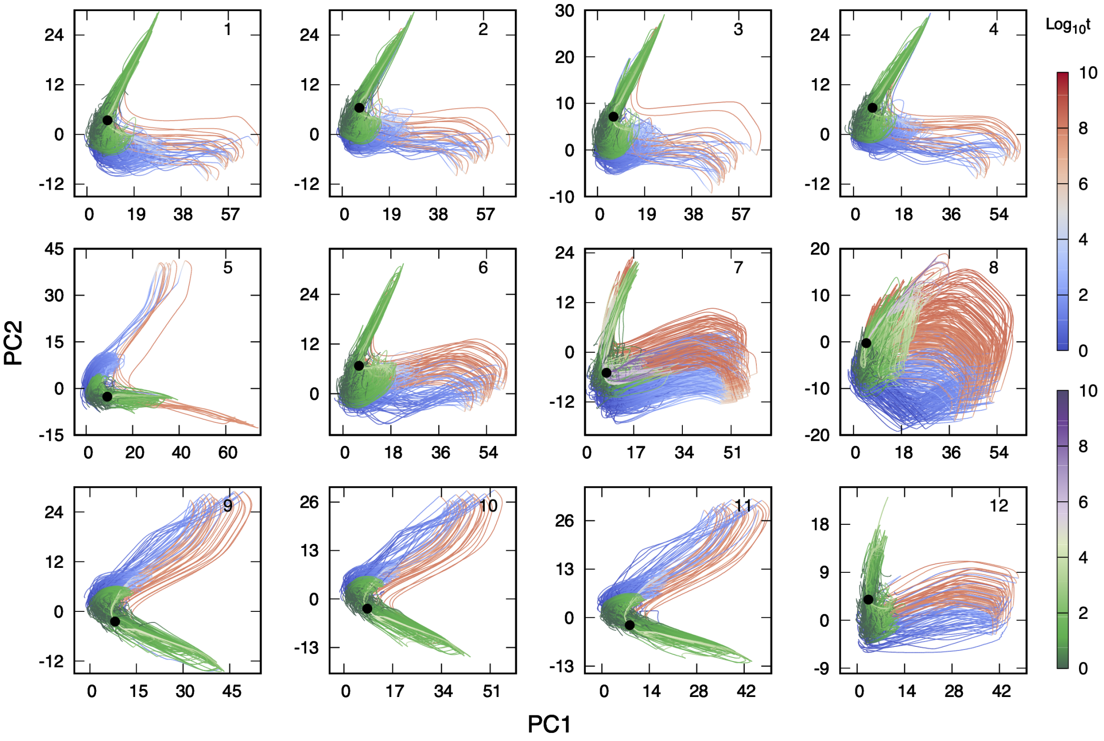

Figure S2: The trajectories on the PCS (from model1 to model12)

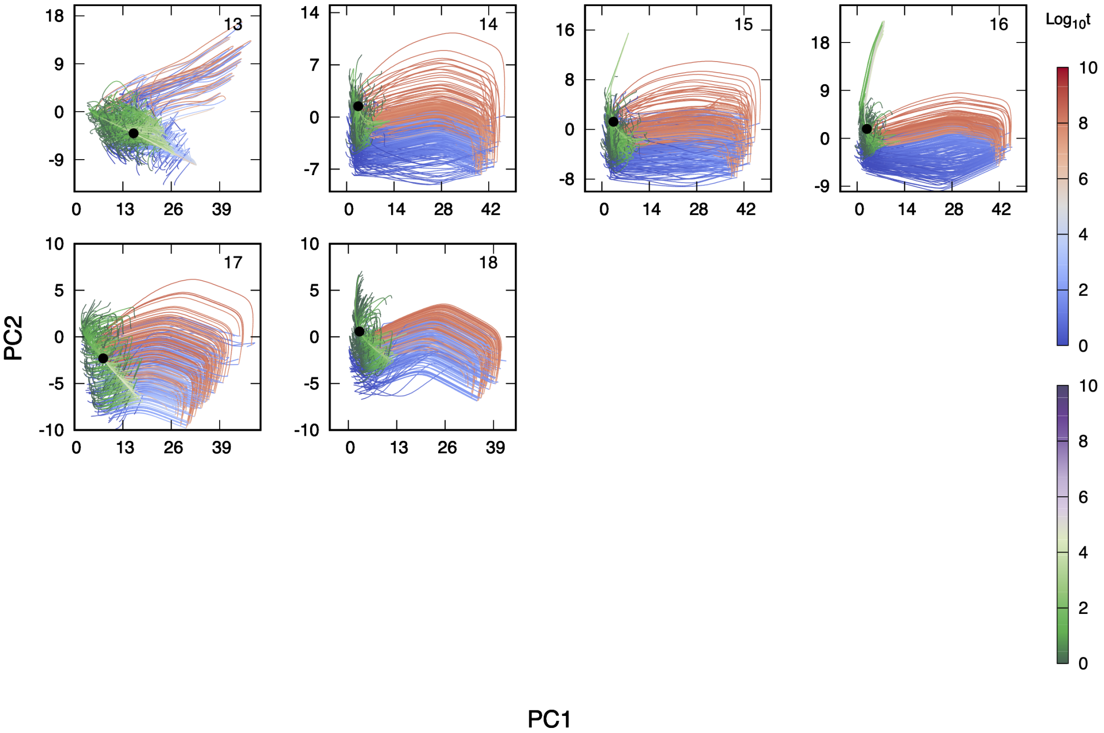

Figure S3: The trajectories on the PCS (from model13 to model18)

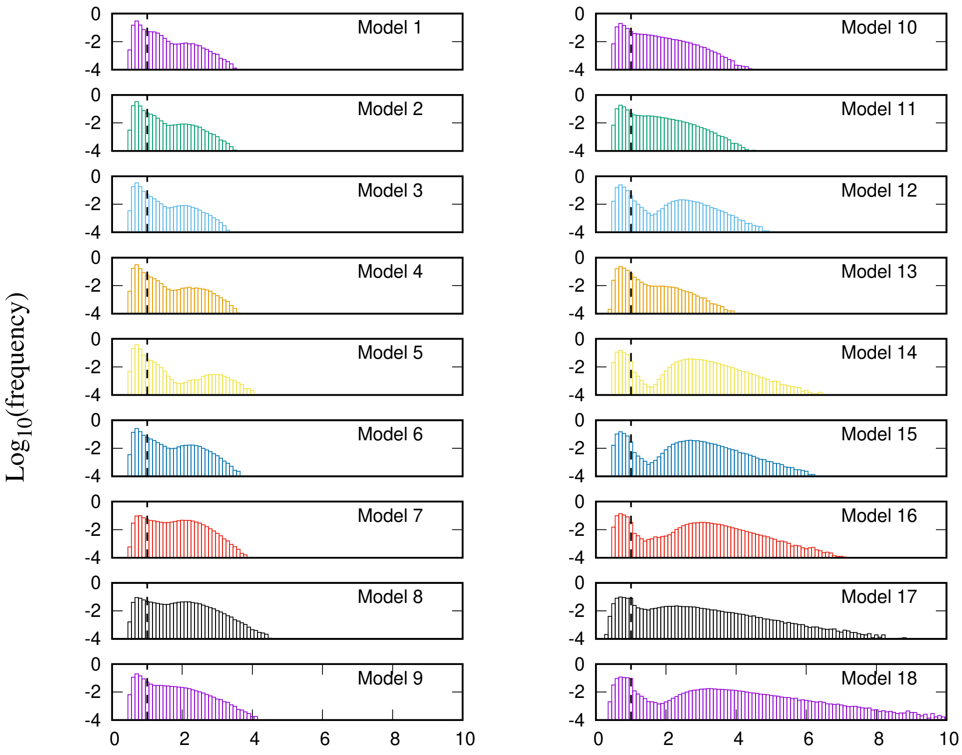

Figure S4: The distribution of the expansion ratio

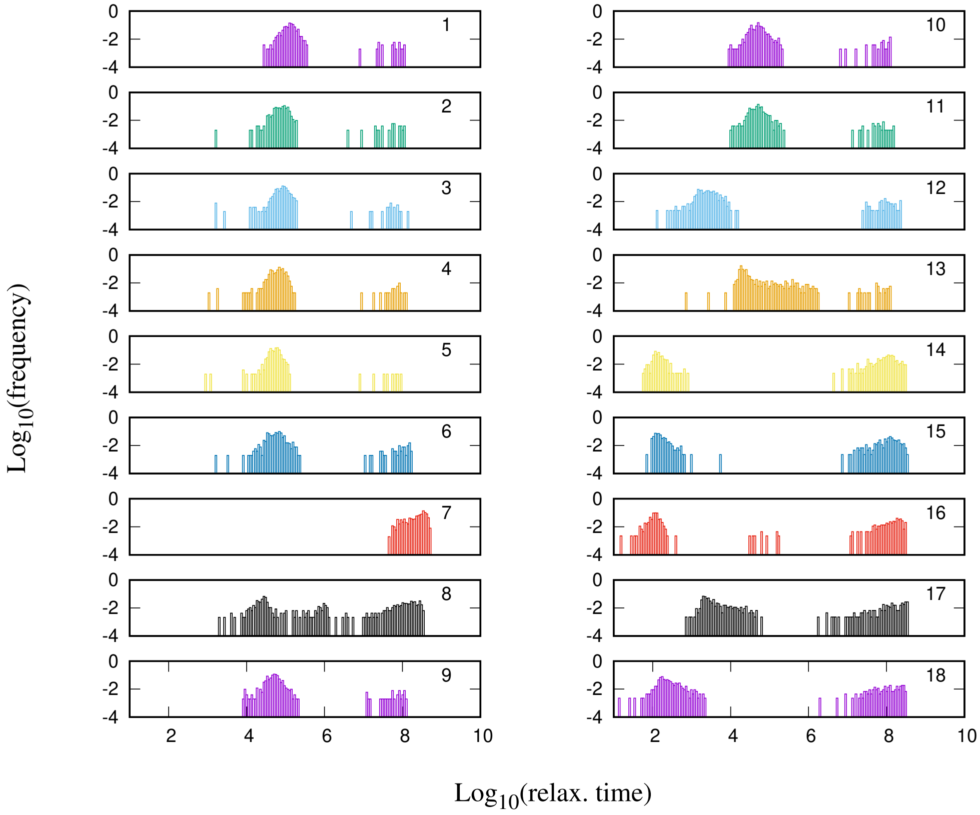

Figure S5: The distribution of the relaxation time

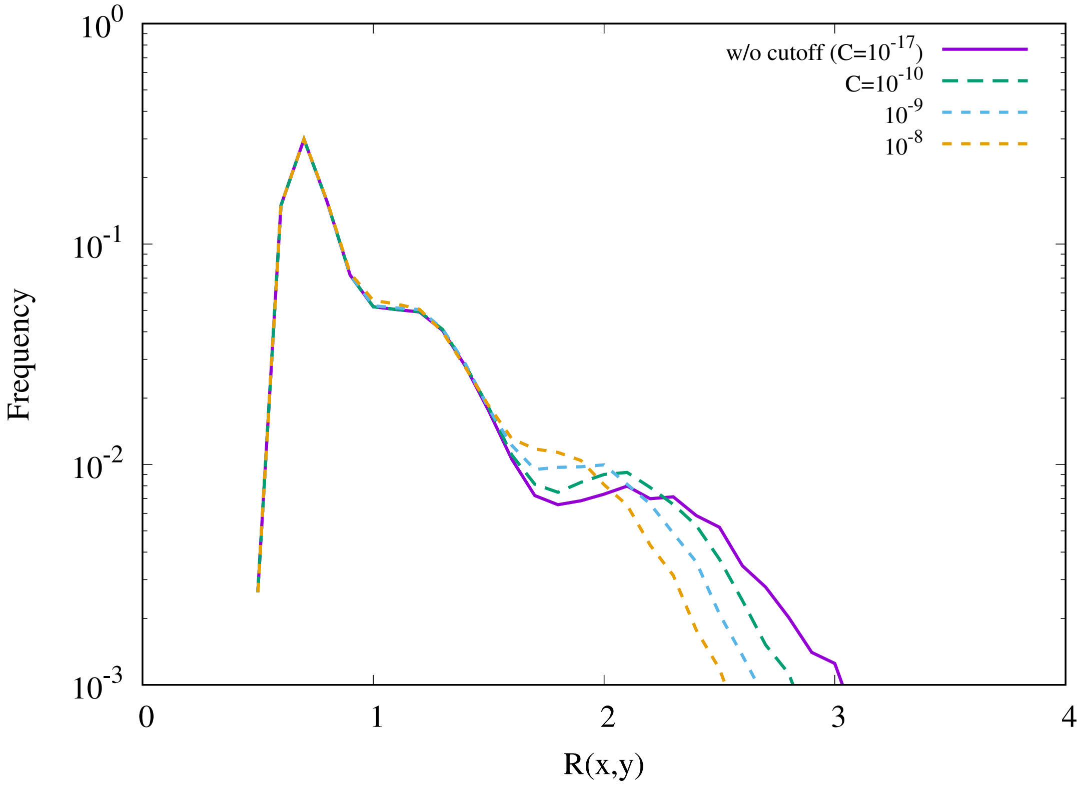

Figure S6: The distribution of the expansion ratio of model0 with several values of cutoff: without cutoff (purple), $C = 10^{-10}$ (green), $C = 10^{-9}$ (cyan), and $C = 10^{-8}$ (orange). Parameters are set to the default values: $v = 1$ and $\kappa = 10^{-6}$ for all reactions, $[\mathrm{glc}] = 1.0, A_t = 1.0, r = 0.1$, and $d = 10^{-8}$.

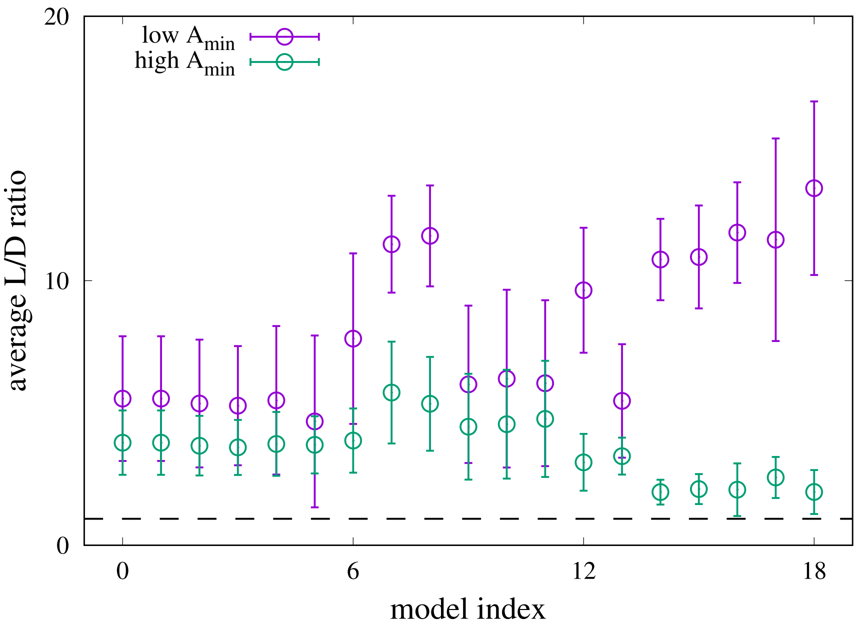

Figure S7: The average ratio of the line integral of the trajectory ($L$) to the Euclidean distance between the initial point and the attractor ($D$) for the growth trajectories and dormant trajectory. The ratio $L/D$ is averaged over the trajectories for each group (high- and low $A_{\min}$ groups) and plotted against the model index with the error bars as the standard deviation. In all the cases, the low $A_{\min}$ trajectories have a larger $L/D$ ratio than the growth trajectories. The broken black line is an eye guideline representing $L/D = 1$.

#### S3.6 The minor attractor of model8

In the model reduction, only model8 exhibited bistability. The fraction of the initial points relaxing to the major attractor, analyzed in the main manuscript, is approximately 92%.

Here, we apply the same analysis for the trajectories relaxing to the minor attractor to confirm that the choice of the attractor is not crucial for model reduction. We applied perturbation on the minor attractor as $10^{u_{i,n}}[X_i^{(\mathrm{ss})}]$ where $u_{i,n}$ and $[X_i^{(\mathrm{ss})}]$ represents a random number for the $i$th metabolite and the $n$th perturbation, uniformly-distributed in $[-1, 1]$ and the concentration of the $i$th metabolite at the minor attractor, respectively.

First, the distribution of the expansion ratio computed from the trajectories relaxing to the minor attractor also exhibited bimodality (Fig. S8A). For the visualization of the trajectories, PCA was performed on the trajectories. In the PC1-PC2 space, the growth trajectories (green-white-purple) and the dormant trajectories (blue-white-red) are clearly separated. Also, the average $L/D$ ratio (see Sec.S3.5) with the standard deviation of the growth- and the dormant trajectories are approximately $7.99 \pm 3.46$ and $9.30 \pm 2.44$, respectively.

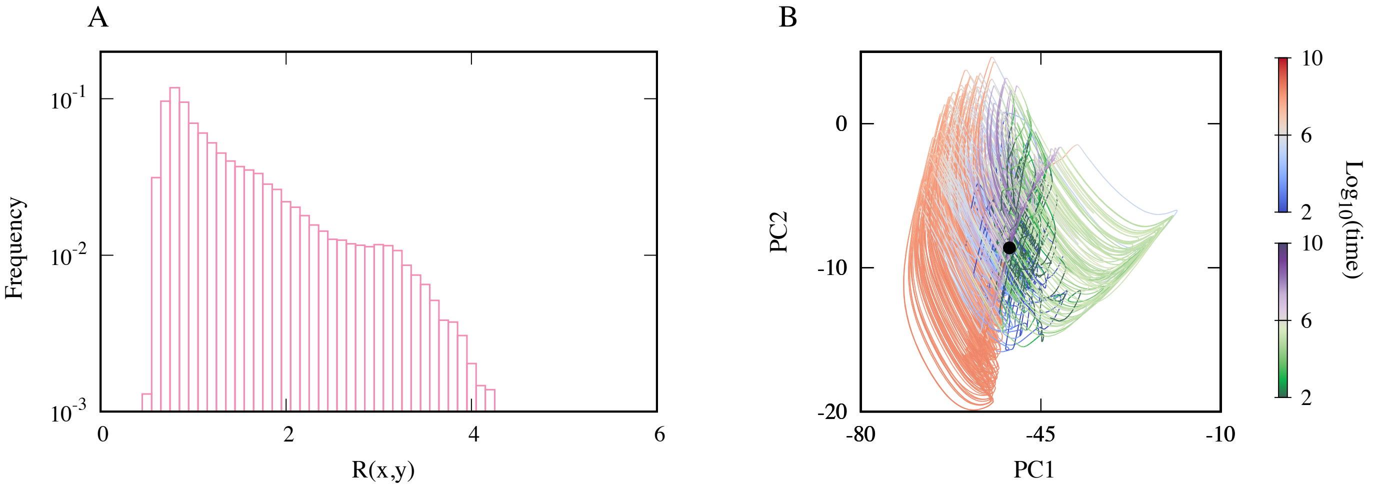

Figure S8: A. The distribution of the expansion ratio of the trajectories perturbed from the minor attractor. B. The trajectories are projected onto the PC1-PC2 space. The trajectories are colored based on the dynamics of $A_{\mathrm{min}}$ (low: blue-white-red, high: green-white-purple). There are 145 growth- and 160 dormant trajectories overlaid in the figure. Parameters are set to the default values: $v = 1$ and $\kappa = 10^{-6}$ for all reactions, $[\mathrm{glc}] = 1.0, A_t = 1.0, r = 0.1$, and $d = 10^{-8}$.

### S4 Analytic solution of the simple model and the choice of the function $\phi$

For obtaining the analytic solution of the simple model (Eqs.(9) and (10) in the main text) in the growth region, we ignore the growth dilution term here ($r = 0$ case). Then, the ordinary differential equation is given by

$$\frac{d[\mathrm{pep}]}{dt} = \phi([\mathrm{pyr}])(1 - [\mathrm{pep}] + [\mathrm{pyr}]) - (1 + d)[\mathrm{pep}],$$
$$\frac{d[\mathrm{pyr}]}{dt} = \phi([\mathrm{pyr}])([\mathrm{pep}] - [\mathrm{pyr}]) - d[\mathrm{pyr}].$$

We solve this set of equations with

$$\phi = \max\{1 - [\mathrm{pyr}], \phi_0\}. \tag{S9}$$

In the region where $\phi([\mathrm{pyr}]) = \phi_0$ holds, the ODE is linear and, thus, easily solved. In the other region, we transform the variables as $\gamma(t) = [\mathrm{pep}](t) + [\mathrm{pyr}](t)$ and $\delta(t) = [\mathrm{pep}](t) - [\mathrm{pyr}](t)$. Then, temporal evolution of $(\gamma, \delta)$ is ruled by

$$\begin{aligned}\frac{d\gamma}{dt} &= 1 - (1+d)\gamma. \\ \frac{d\delta}{dt} &= (1 - (\gamma - \delta)/2)(1 - 2\delta) - (\gamma + \delta)/2 - d\delta.\end{aligned}$$

The solution for this is given by

$$\begin{aligned}\gamma(t) &= f^{-1} + C_0 e^{-ft}, \\ \delta(t) &= C_0 e^{-ft} - (\kappa + \eta)/2f, \\ &+ C_0 \frac{\alpha C_1 U(1+\alpha, 2+\beta; \zeta e^{-ft}) + L^{1+\beta}_{-(1+\alpha)}(\zeta e^{-ft})}{C_1 U(\alpha, 1+\beta; \zeta e^{-ft}) + L^{\beta}_{-\alpha}(\zeta e^{-ft})} e^{-ft}.\end{aligned} \tag{S10}$$

where $U$ and $L$ are the confluent hypergeometric function and the associated Laguerre polynomial, respectively. Lumped parameters are

$$\begin{aligned}f &= 1 + d, \\ \xi(f) &= 1 - 3f + f^2, \\ \eta(f) &= \sqrt{1 - 6f + 3f^2 + 2f^3 + f^4}, \\ \zeta(f) &= \frac{C_0}{f}, \\ \kappa(f) &= -1 + f + f^2, \\ \alpha(f) &= (\xi(f) + \eta(f))/2f^2, \\ \beta(f) &= \eta(f)/f^2,\end{aligned}$$

and the integral constants $C_0$ and $C_1$ are given by

$$\begin{aligned}C_0 &= \gamma(0) - 1/f, \\ C_1 &= -\frac{C_0 L^{1+\beta}_{-(1+\alpha)}(C_0/f) + L^{\beta}_{-\alpha}(C_0/f)\Big(C_0 - (\kappa+\eta)/2f - \delta(0)\Big)}{C_0 \alpha U(1+\alpha, 2+\beta; C_0/f) + U(\alpha, 1+\beta; C_0/f)\Big(C_0 - (\kappa+\eta)/2f - \delta(0)\Big)}.\end{aligned}$$

Note that there is only a single timescale $1/f = 1/(1+d)$ in the growth region. $1/f$ is $\mathcal{O}(1)$ with the default parameter set. While we omitted the growth-dilution term $-\mu[\cdot]$ for obtaining the analytic solution, if the growth rate $\mu$ is smaller than 1, the effect of including the dilution term is masked by $f$. On the other hand, it simply speeds up the relaxation if $\mu$ is larger than unity. Thus, the inclusion of $\mu$ does not change the argument that the slowest timescale in the growth region is $\mathcal{O}(1)$.

The analytic solution is obtained for a specific functional form of $\phi$ defined as eq. (S9). However, the structure of the vector field is not sensitive to the choice of $\phi$. In Figs. S9, we drew the two-dimensional vector fields with the functional form of $\phi$ chosen to be an exponential function $\phi = \exp(-[\mathrm{pyr}])$(A) and a Hill function $\phi = [\mathrm{pyr}]^{n_H}/(K^{n_H} + [\mathrm{pyr}]^{n_H})$ (B). The figures imply that the nature of the vector field is robust as long as $\phi$ reaches a small value as [pyr] increases.

### S5 The minimal model with de-novo AMP synthesis

Since the *E. coli* core model includes no AMP synthesis pathway, we assumed that the total concentration of the adenine nucleotide carriers (ATP, ADP, and AMP) is constant in the main text. To check if this

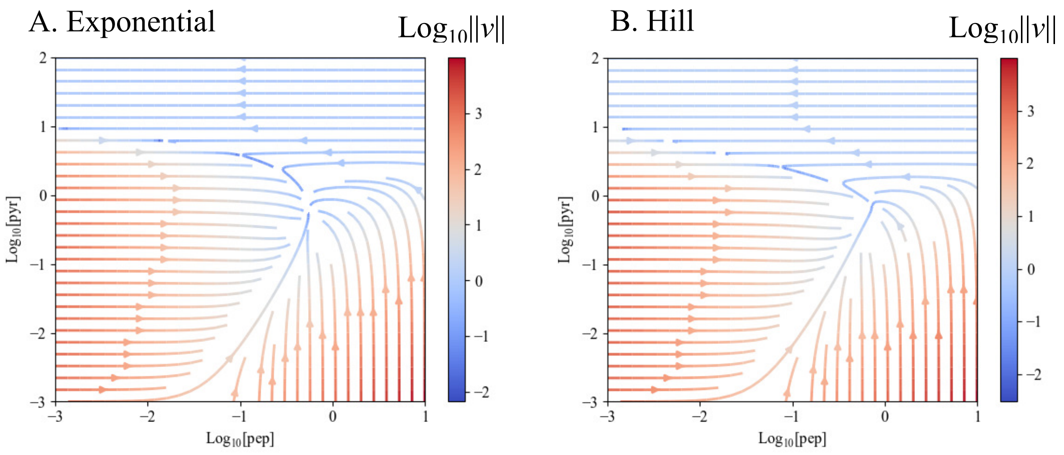

Figure S9: The streamlines in the phase space with alternative functions. The exponential function $\exp(-[\mathrm{pyr}])$ and the Hill function $[\mathrm{pyr}]^{n_H}/(K^{n_H} + [\mathrm{pyr}]^{n_H})$ are used as the function $\phi$ for A and B, respectively. $K = 1.0$ and $n_H = -4$ for B. The other parameter values are $v = 1$ and $\kappa = 10^{-6}$ for all reactions, $[\mathrm{glc}] = 1.0, r = 0.1$, and $d = 10^{-8}$.

assumption is crucial for the obtained result, we introduce a coarse-grained AMP synthesis reaction to the minimal model and study the dynamics of the model.

Here, we extend the minimal model studied in the main manuscript. The nucleotide carriers such as AMP and GMP are synthesized from the pentose-phosphate pathway (PPP) chemicals by utilizing ATP energy. In the minimal model, PPP is already removed from the model; thus, glucose is the chemical species closest to the PPP in the original metabolic network. Therefore, we introduced phenomenological AMP synthesis reaction glc+atp $\rightleftharpoons$ amp+adp where glucose is the substrate, and the reaction needs the energy consumption (ATP $\rightarrow$ ADP). Then, the total concentration of adenine carriers is no longer constant, and thus, we put the constant-rate degradation term and the growth dilution term whose rate is proportional to the growth reaction to all chemicals. Then, the equations are given by

$$\frac{d[\mathrm{pep}]}{dt} = J_{\mathrm{uptake}} + J_{\mathrm{pps}} - J_{\mathrm{pyk}} - J_{\mathrm{ppc}} - (d+\mu)[\mathrm{pep}], \tag{S11}$$

$$\frac{d[\mathrm{pyr}]}{dt} = J_{\mathrm{pyk}} - J_{\mathrm{pps}} - (d+\mu)[\mathrm{pyr}], \tag{S12}$$

$$\frac{d[\mathrm{oaa}]}{dt} = J_{\mathrm{pps}} - J_{\mathrm{growth}} - (d+\mu)[\mathrm{oaa}], \tag{S13}$$

$$\frac{d[\mathrm{atp}]}{dt} = J_{\mathrm{uptake}} + J_{\mathrm{pyk}} - J_{\mathrm{pps}} - J_{\mathrm{growth}} - J_{\mathrm{adk1}} - J_{\mathrm{amps}} - (d+\mu)[\mathrm{atp}], \tag{S14}$$

$$\frac{d[\mathrm{adp}]}{dt} = -J_{\mathrm{uptake}} - J_{\mathrm{pyk}} + J_{\mathrm{growth}} + 2J_{\mathrm{adk1}} + J_{\mathrm{amps}} - (d+\mu)[\mathrm{adp}], \tag{S15}$$

$$\frac{d[\mathrm{amp}]}{dt} = J_{\mathrm{pps}} - J_{\mathrm{akd1}} + J_{\mathrm{amps}} - (d+\mu)[\mathrm{amp}], \tag{S16}$$

$$J_{\mathrm{amps}} = v_{\mathrm{atp}}([\mathrm{glc}][\mathrm{atp}] - [\mathrm{amp}][\mathrm{adp}]). \tag{S17}$$

Here we analyzed the trajectories starting from randomly-generated initial point $10^{u_{i,n}}$ with $u_{i,n}$ as the uniformly-distributed random number in $[-1, 1]$ for the $i$th chemical and the $n$th initial point. This reduces the requirements of the computational resources because we can skip the computation for finding attractors. As far as we have checked, the model had a single attractor.

Fig. S10 shows the distribution of the expansion ratio and the projected trajectories onto the two-dimensional PC space. As depicted, the model with the de-novo synthesis of AMP still exhibits distinct trajectories while the dormant trajectories become rare with the default parameter set.

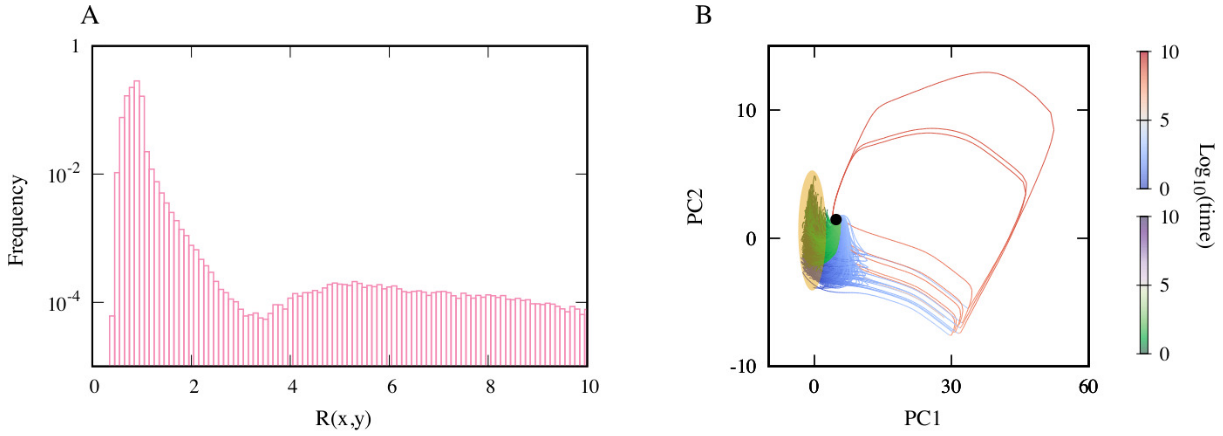

Figure S10: A. The distribution of the expansion ratio of the minimal model with the de-novo AMP synthesis. It shows clear bimodality. B. The trajectories projected onto the 2-dimensional PC space. The green-white-purple and blue-white-red colored trajectories are the growth- and the dormant trajectories, respectively. The trajectories are colored based on the relaxation time of each trajectory. The black dot represents the attractor and the initial points cluster in the region highlighted in orange. $v = 1$ and $\kappa = 10^{-6}$ for all reactions while $v_{\text{atp}} = 0.1$, $[\text{glc}] = 1.0, r = 0.1$, and $d = 10^{-8}$).

### S6 model0 with the nicotinamide nucleotide carriers

In the main text, we replaced NAD(NADP) and NADH(NADPH) with ATP and ADP, respectively, with the assumption that the ATP synthesis via the electron transport chain and the conversion of NADP to NADPH is sufficiently quick. Here, we relax these assumptions and introduce the dynamics of NAD, NADH, NADP, and NADPH to model0.

Here, we introduce two phenomenological reactions shown in Table.1, and the replacement of the nicotinamide nucleotide carriers by the adenine nucleotide carriers is not performed. A full list of the reactions is provided in *SI Data*.3.

| Reaction Name | Reaction Formula |
|---|---|
| ATPPMF | NADH + ADP → NAD + ATP |
| NADTRHD | NAD + NADPH → NADH + NADP |

Table 1: Reactions added to model0

The reaction "ATPPMF" is for the ATP generation using proton motive force which consists of NADH16, CYTBD, and ATPS4r in the original core model. NADTRHD has the same stoichiometry as that in the core model except for the hydrogen ion.

In the model, the degradation and growth-dilution term are omitted for their dynamics, and [nad]+[nadh] and [nadp]+[nadph] are constant because the cofactors are not newly synthesized in the model. For simplicity, here we set $[\text{atp}] + [\text{adp}] + [\text{amp}] = [\text{nad}] + [\text{nadh}] = [\text{nadp}] + [\text{nadph}] = A_t$.

Here we used the randomly-generated initial conditions with $u_{i,n}$ as the random number, $10^{u_{i,n}}, u_{i,n} \in [-1, 1]$, instead of the initial condition generated by the perturbation. The concentrations of the cofactors are normalized after assigning the random numbers. We found in this model the distinct trajectories emerge when the range of initial concentration of pyruvate is set to $[1, 10^3]$ (i.e, $u_{i,n} \in [0, 3]$ for pyruvate) as shown in

Figs. S11. This is qualitatively consistent with the result of the minimal model in the main text that pyruvate plays a crucial role in displaying distinct relaxation behaviors. Including the nicotinamide nucleotide carries simply changes the needed pyruvate level to have a dormant trajectory quantitatively.

In this model, the separation of the trajectories is unclear in the two-dimensional PC space (Fig. S11B), while it is in the three-dimensional PC space (Fig. S11C). Note that, in these figures, we colored the trajectories based on $A_{\min}$ because the distribution of $A_{\min}$ showed clear bimodality. However, it is not fully consistent with the separation of the trajectories in Fig. S11C. This is probably because $A_{\min}$ alone is now an insufficient indicator of the energetic state of the cell. For more precise coloring, the contributions of the nicotinamide nucleotide carriers should be incorporated, while it is beyond the scope of the present study.

### S7 Model reduction with random order

In the main text, We have obtained the minimal model by manually deciding the order of the reaction removal. However, in general, the resulting minimal models by the reduction depend on the order of the removal. Here, we reduce the *E. coli* core model in random order to investigate if the obtained minimal models share features in common with the ones that summarise the l presented in the main text.

Pseudo-codes that summarise the algorithms are presented in the following. For the random reduction, we randomly choose a reaction and check if the reaction is removable by Algorithm S1 and iterate it until there is no more removable reaction.

Algorithm S1 requires the reaction network (a list of the reactions) and the name of the reaction to be removed as inputs. If the reaction is removable from the network, it returns the reaction network without the reaction, while it returns the same reaction network as the input if the reaction is not removable.

The algorithm first checks whether the input reaction can be simply removed (line 3) or the contraction is needed (line 6). In the case where the removal of the input reaction leads to dead-end chemicals (chemicals with only one reaction connected), the algorithm computes a set of reactions $T$. $T$ is a minimal set of reactions, including the input reaction so the simultaneous removal of the reactions in $T$ from the reaction network does not lead to dead-end chemicals (line 10).

If the reaction removal eliminates a chemical in the growth reaction, a neighboring chemical in the backbone network $B$ (the reaction network without ATP, ADP, and AMP) is chosen as the replacement of the eliminated chemical (line $17-19$).

By removing the reaction, we obtain a candidate of the reduced reaction network $\tilde{R}$. Then, the algorithm checks if the network $\tilde{R}$ satisfies the following three conditions: connectivity, a non-zero steady flux without the degradation and growth dilution, and multimodality of the distribution of the expansion ratio (line 20).

Algorithm S2 calls Algorithm S1 with a randomly selected reaction(s) and checks if the obtained network is minimal.

To check if the distribution of the expansion ratio $\tilde{R}$ is multimodal and/or long-tailed, we calculated $\tilde{R}$ of the kinetic model with a default parameter set used in the main text ($v = 1$ and $\kappa = 10^{-6}$ for all reactions.

The reaction lists of 16 minimal models obtained from different random seeds for the reduction are given in *SI Data*.4. By random reduction, we obtained two groups of minimal models classified by the shape of the distribution of the expansion ratio. The first case shows clear multimodality (model $\#0-\#13$). The second case shows a long-tail rather than additional peaks (model #14 and #15).

All the minimal models had more reactions than the minimal model in the main text. Interestingly, the network structures of the minimal models are qualitatively different depending on whether the model exhibits the clear bimodal distribution of the expansion ratio or not. The models with the bimodal distribution share two network features, namely, (i) ATP, ADP, and AMP are in the model, and (ii) there are both types of reactions; with- and without- the adenine nucleotide carriers coupling as well as branching of the network. These are the vital network features for the model to satisfy the two conditions for the distinct trajectories discussed in the main text. We like to emphasize that during the random model reductions, several models without AMP were generated, while none of them showed a multimodal distribution of the expansion ratio, and they were rejected based on the distribution of the expansion ratio.

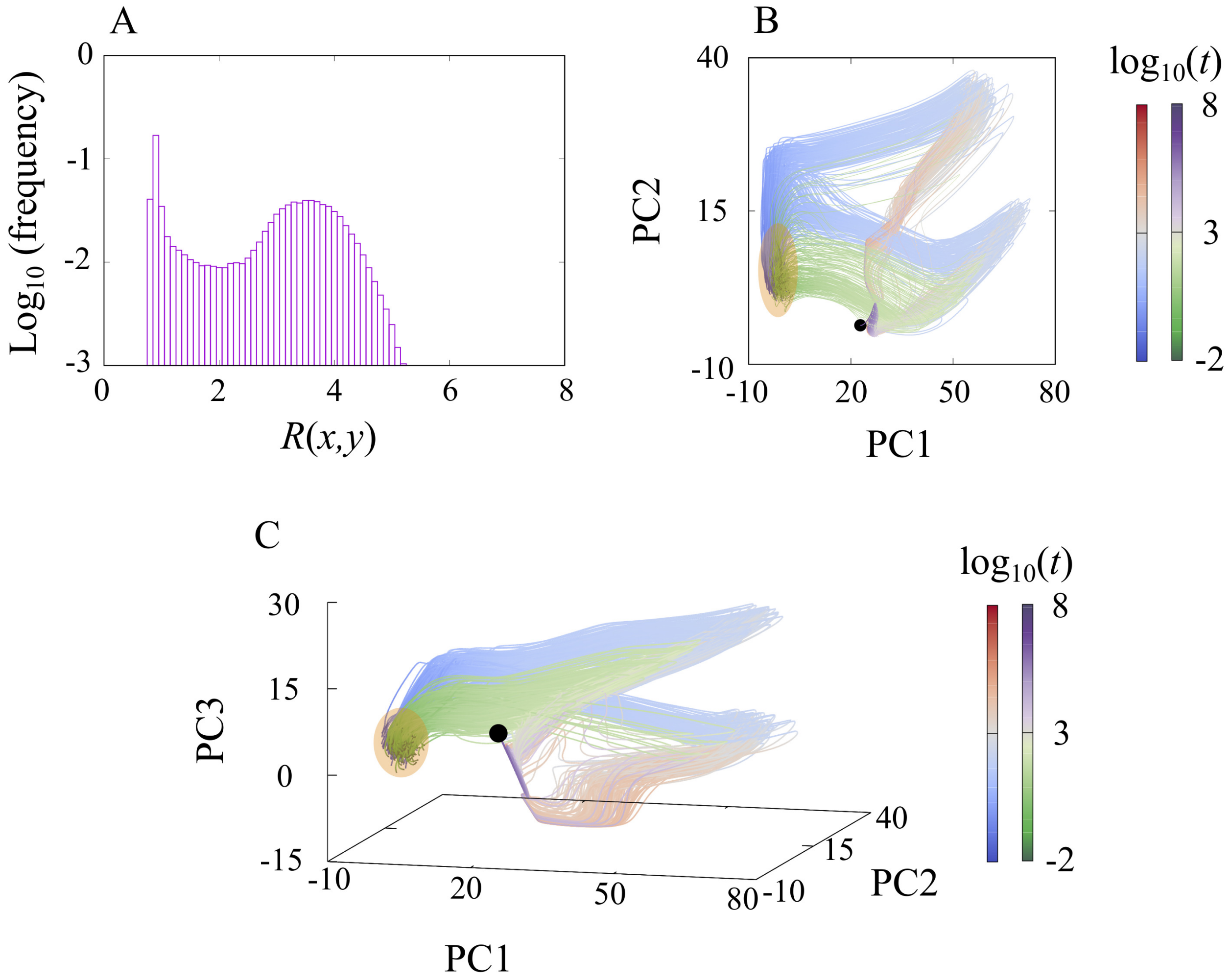

Figure S11: A. The distribution of the expansion ratio of the model0 with NAD, NADH, NADP, and NADPH, showing clear bimodality. B and C. The trajectories in the 2-dimensional (B) and 3-dimensional (C) PC space. The green-white-purple and blue-white-red colored trajectories are the growth- and the dormant trajectories, respectively. The trajectories are colored based on $A_{\min}$. The black dot represents the attractor and the initial points cluster in the region highlighted in orange. $v = 1$ and $\kappa = 10^{-6}$ for all reactions. Other parameters are $[\mathrm{glc}] = 1.0, A_t = 1/3, r = 0.1$, and $d = 10^{-8}$.

**Algorithm S1** Compute a reduced network from given network $R$ and a reaction to be removed `rxn`

Notations:

- $R$ : the reaction network
- $B$ : the reaction network without atp, adp, and amp
- $R - T$ : the reaction network without the reactions in $T$
- $E(k)$ : the number of reactions that the chemical $k$ is associated

$C \leftarrow$ the list of chemicals in $R$
$C_0 \leftarrow C-$`'glc'`
**if** $E(k) \geq 2$ for $^{\forall}k \in C_0$ after `rxn` is removed **then**
  `RemoveList`$\leftarrow$[`rxn`]
  `RenameList`$\leftarrow$[]
**else if** `rxn` is one-to-one reaction in $B$ **and** no loop b/w substrate and product of `rxn` **then**
  `RemoveList`$\leftarrow$[`rxn`]
  `RenameList`$\leftarrow$[(Substrate of `rxn`,Product of `rxn`)]
**else**
  find a minimal reaction set $T$ so that $E(k) \geq 2$ or $E(k) = 0$ for $^{\forall}k \in C_0$ in $R - T$
  `RemoveList`$\leftarrow T$
  `RenameList`$\leftarrow$[]
**end if**
$\tilde{R} \leftarrow R-$`RemoveList`
$\tilde{C} \leftarrow$chemicals in $\tilde{R}$
Rename chemical names in $\tilde{R}$ and $\tilde{C}$ according to `RenameList`
**if** a growth factor $g_i$ is removed **then**
  replace $g_i$ by a closest chemical on $B$
**end if**
**if** Connected **and** Non-zero steady flux exists **and** The dist. of the exp. ratio is multimodal **then**
  **return** $\tilde{R}$
**else**
  **return** $R$
**end if**

On the other hand, the minimal models exhibiting the distribution of the expansion ratio with a long-tail lack the second condition, i.e., all the reactions are coupled with the adenine nucleotide carriers. As a consequence, all the metabolic reactions are uniformly slowed down even if ATP and ADP deplete. Thus, the distinction among the trajectories is not as clear as in the other minimal models.

In Figs. S13, we plotted the trajectories of each minimal model. Since we found that the distributions of $A_{\min}$ (see Section S3.3) of the minimal models were double-peaked, we colored the trajectories by the same criterion used in Section S3.3. Interestingly, there are several types of minimal models in terms of the visual impression of the trajectories in PC1-PC2 space; The minimal models show clear separations of two types of trajectories as the minimal model studied in the main text ($\#1, \#2, \#6 - \#8, \#11$ and $\#12$), models exhibiting the oscillation during the relaxation ($\#3$ and $\#10$), and models where the separation of the trajectories is not quite clear ($\#0, \#4, \#5, \#9$ and $\#13$)[1]. However, in the original high-dimensional phase-space, the two types of trajectories are distinct in terms of the $L/D$ ratio (see Section S3.5) for the models with the bimodal distribution of the expansion ratio (model #0-#13) as shown in Fig. S14.

**Algorithm S2** The algorithm for a random reduction (the same notation with Algorithm S1 is used)

**while** 1 **do**
  `RxnList` $\leftarrow$ All reactions in $R-$['growth reaction']
  Shuffle `RxnList`
  **for** r in `RxnList` **do**
    $R_0 \leftarrow$SingleLoopReduction($R$,$r$) (*see Alg. S1*)
    **if** $R_0 \neq R$ **then**
      break
    **end if**
  **end for**
  **if** $R = R_0$ **then**
    **return** R
  **end if**
  $R \leftarrow R_0$
**end while**

### S8 The behavior of model0 with randomly assigned parameters

In the main text, we saw that the distinct trajectories emerge in two sets of parameter values, the realistic setting and uniform assignment for $v_i'$s and $k_i'$s. Here, we check the robustness of the emergence of the distinct trajectories by randomly assigning the parameter values.

We simulated model0 with a variety of parameter values. As concluded in the main text, the concentrations of ATP and ADP play a crucial role in the emergence of distinct trajectories. Therefore, we studied the relaxation dynamics of the model with several values of the total concentrations of the adenine nucleotide carriers $A_t$ (= [atp] + [adp] + [amp]). In addition, we assigned random values for the kinetic parameters in the rate equations ($v_i'$s and $k_i'$s. see Eq.(3) in the main text). We kept the concentration of the nutrient [glc], the degradation constant $d$, and the proportionality constant between the growth reaction and the growth rate $r$ the same as the main text.

For each values of $A_t$, we generated 32 random vectors of parameters $\vec{p} = (\vec{v}, \vec{k})$ where $\vec{v}$ and $\vec{k}$ are vector representation of the parameters $v_i'$s and $k_i'$s, respectively. For each $\vec{p}$, we ran the differential equations from 128 randomly generated initial points and computed the distribution of the expansion ratio. $v_i'$s and $k_i'$s are given as $10^u$ where $u$ is an uniformly-distributed random number. For $v_i'$s, $u$ ranges from 0 to 1, while it ranges from $-6$ to $-4$ or from $-4$ to $-2$ for $k_i'$s.

[1]Note that the reductions were made in random order and the same minimal network can result. Actually, there are several the same model pairs, namely, #0 and #13, #1 and #7, #3 and #10 and #4 and #9.

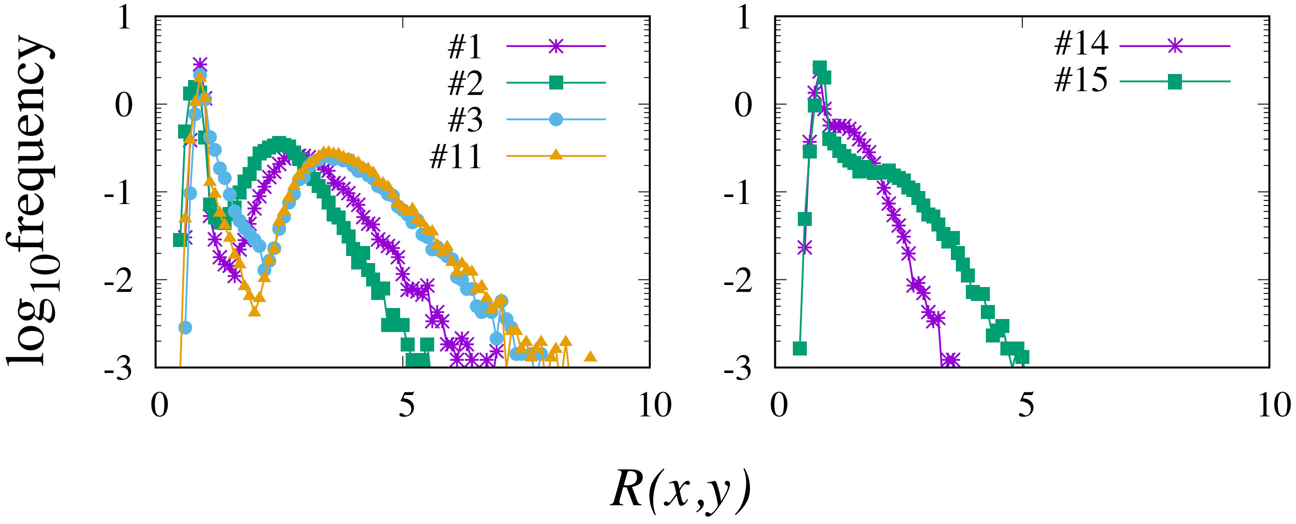

Figure S12: Two types of resulted distribution of the expansion ratio. The first type exhibiting the clear multimodality (left) and the second type showing a long-tail rather than additional extrema (right). Labels in the panels are the indices of the minimal models.

Fig. S15 shows the fraction of $\vec{p}$'s that led a bimodal distribution as a function of $A_t$. The results obtained from two different ranges of $k_i'$s are overlaid. The bimodality is judged by using the same criterion described in Sec.S3.2. The fraction of parameter sets leading to a bimodal distribution of the expansion ratio is a decreasing function of $A_t$ if $k_i'$s ranges from $10^{-4}$ to $10^{-2}$, while interestingly, it shows non-monotonic behaviour in the case where $k_i'$s ranges from $10^{-6}$ to $10^{-4}$. Thus, the emergence of distinct trajectories robustly takes place while the chance of it with random parameter assignments eventually decreases as $A_t$ increases.

### References

[1] Yikun Tan and James C Liao. Metabolic ensemble modeling for strain engineers. *Biotechnology journal*, 7(3):343–353, 2012.

[2] Ali Khodayari, Ali R Zomorrodi, James C Liao, and Costas D Maranas. A kinetic model of escherichia coli core metabolism satisfying multiple sets of mutant flux data. *Metabolic engineering*, 25:50–62, 2014.

[3] Christopher M Bishop. *Pattern recognition and machine learning*. springer, 2006.

[4] Luca Gerosa, Bart RB Haverkorn van Rijsewijk, Dimitris Christodoulou, Karl Kochanowski, Thomas SB Schmidt, Elad Noor, and Uwe Sauer. Pseudo-transition analysis identifies the key regulators of dynamic metabolic adaptations from steady-state data. *Cell systems*, 1(4):270–282, 2015.

[5] Amir Akbari, James T Yurkovich, Daniel C Zielinski, and Bernhard O Palsson. The quantitative metabolome is shaped by abiotic constraints. *Nature Communications*, 12(1):1–19, 2021.

[6] Athel Cornish-Bowden. *Fundamentals of enzyme kinetics*. John Wiley & Sons, 2013.

[7] F. Pedregosa, G. Varoquaux, A. Gramfort, V. Michel, B. Thirion, O. Grisel, M. Blondel, P. Prettenhofer, R. Weiss, V. Dubourg, J. Vanderplas, A. Passos, D. Cournapeau, M. Brucher, M. Perrot, and E. Duchesnay. Scikit-learn: Machine learning in Python. *Journal of Machine Learning Research*, 12:2825–2830, 2011.

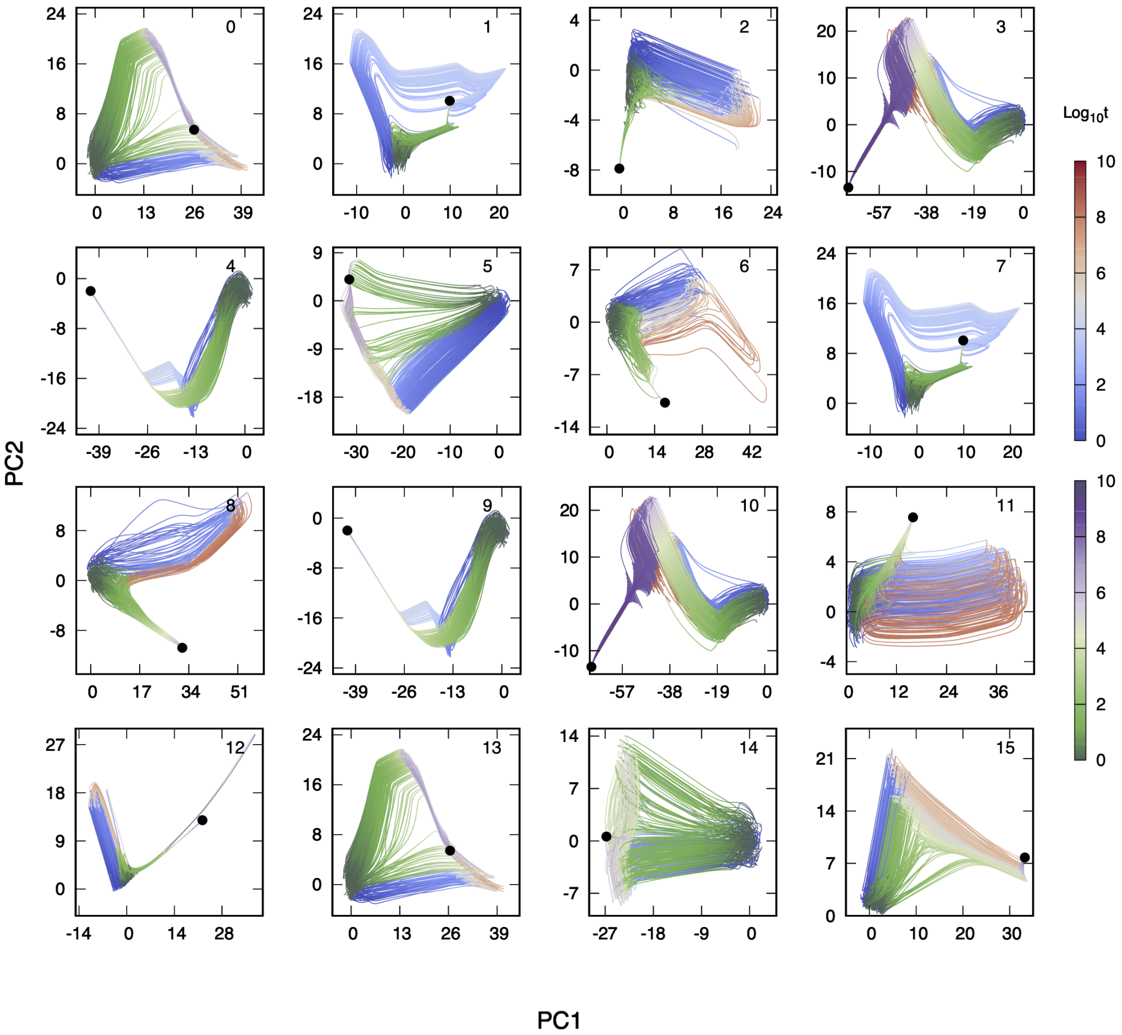

Figure S13: The trajectories on the PCS. Trajectories are colored according to $A_{\min}$.

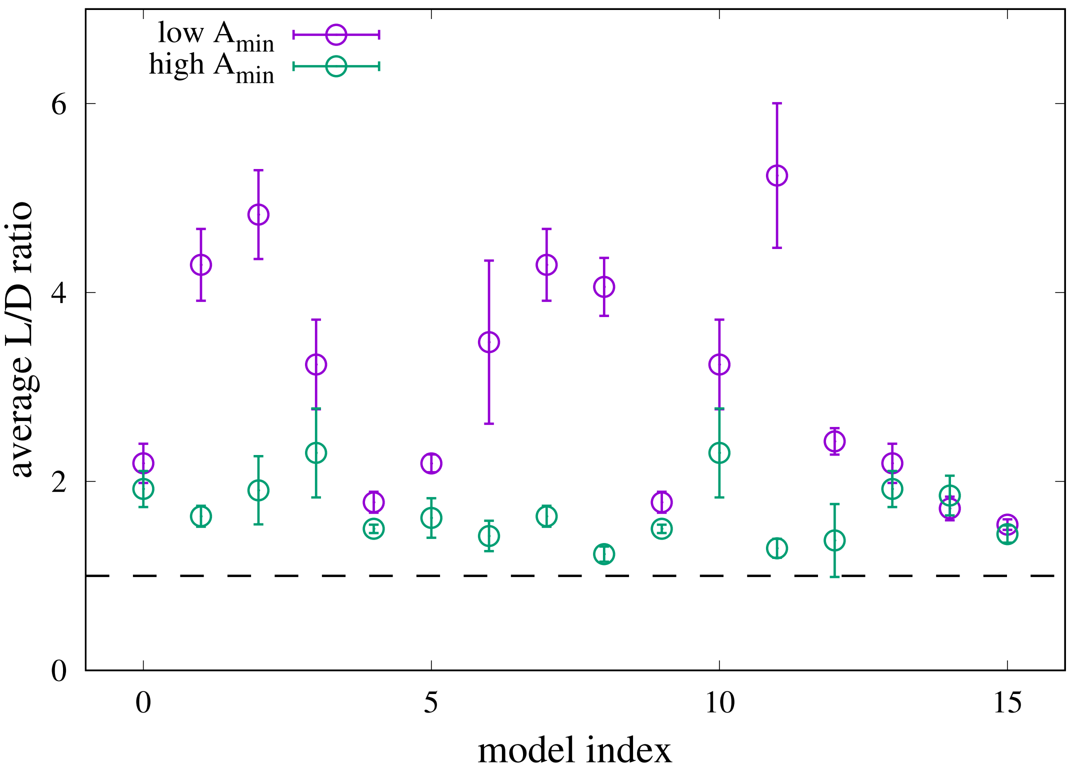

Figure S14: The average $L/D$ ratio of for the minimal models obtained by the random reduction. Error bars indicate the standard deviation. The trajectories with low $A_{\min}$ has a higher $L/D$ ratio than that of trajectories with high $A_{\min}$. The black dashed line is $L/D = 1$ for an eye guide.

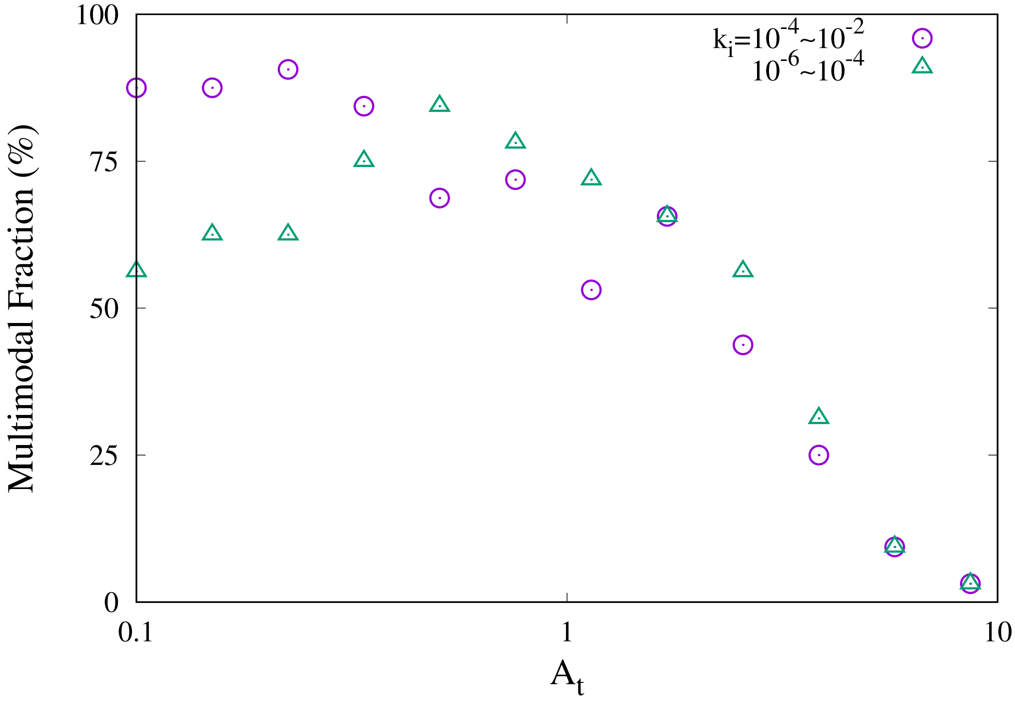

Figure S15: The fraction of the parameter sets leading to a multimodal distribution of the expansion ratio is plotted as the function of the total adenine nucleotide carriers concentration, $A_t$. The result obtained from the simulations with two different ranges of $k_i'$s are overlaid.
